# Beyond polyA: scalable single-cell total RNA-seq unifies coding and non-coding transcriptomics

**DOI:** 10.1101/2025.08.08.669394

**Authors:** Alina Isakova, Daniel Dan Liu, Ivana Cvijovic, Rahul Sinha, Anna E Eastman, Sirle Saul, Angela Detweiler, Norma Neff, Shirit Einav, Irving L Weissman, Stephen R. Quake

**Author notes:** Correspondence: Stephen R. Quake, Alina Isakova.

## Abstract

Non-coding RNAs represent a widespread and diverse layer of post-transcriptional regulation across cell types and states, yet much of their diversity remains uncharted at single-cell resolution. This gap stems from the limitations of widely used single-cell RNA-sequencing protocols, which focus on polyadenylated transcripts and miss many short or non-polyadenylated RNAs. Here, we adapted single-cell RNA-sequencing on the 10x Genomics platform to capture a broad complement of coding and non-coding RNAs—including miRNAs, tRNAs, lncRNAs, histone RNAs, and non-adenylated viral transcripts. This approach enabled the discovery of rich, dynamic non-coding RNA programs across immune cells, virally infected hepatocytes, and the developing human brain. In dengue virus-infected hepatocytes, we detect non-adenylated viral transcripts and distinguish active from transcriptionally quiescent infected states, each with distinct host regulatory signatures. In brain tissue, we identify biotype-specific, cell-type– restricted non-coding RNAs, including miRNAs whose expression anticorrelates with predicted targets, consistent with post-transcriptional regulatory relationships. We show that *MIR137*, one of the strongest GWAS loci associated with schizophrenia and intellectual disability, is expressed specifically in Cajal-Retzius cells, an early-born but transient population that guides subsequent cortical neuron migration. These findings demonstrate the importance of non-coding RNAs in defining cell identity and state, and show how expanded transcriptome coverage can reveal additional layers of gene control—now accessible through practical and scalable single-cell profiling.

## Introduction

Non-polyadenylated RNAs comprise a substantial and functionally diverse portion of the transcriptome. These include microRNAs (miRNAs), long non-coding RNAs (lncRNAs), small nucleolar and small nuclear RNAs (snoRNAs, snRNAs), circular RNAs (circRNAs), transfer RNAs (tRNAs), histone mRNAs^1,2^, and many viral transcripts that naturally lack polyA tails^3,4^. Together, they play essential roles in transcriptional regulation, RNA processing, translation, stress response, and cell fate transitions^5–8^. Yet, these transcripts remain largely invisible to conventional single-cell RNA sequencing (scRNA-seq) platforms, which rely on polyA-capture and are consequently biased toward protein-coding mRNAs^9^.

While specialized protocols such as Smart-seq-total^10^, VASA-seq^11^, RamDA-seq^12^, snapTotal-seq^13^, MATQ-seq^14^, scComplete-seq^15^ have expanded single-cell profiling to include non-polyadenylated RNAs, these methods often require custom equipment, custom enzymes, extensive sample processing, or bespoke computational pipelines—factors that limit their scalability and integration into high-throughput and widely-adopted workflows. Moreover, most of these approaches either underperform or entirely fail to capture mature microRNAs, which play critical roles in development, immunity, and disease. As a result, large portions of RNA biology are systematically excluded from most single-cell studies, leaving blind spots in efforts to map regulatory networks, cell states, and disease mechanisms which involve non-coding transcripts.

Here, we present a generalizable and scalable framework for total RNA profiling in single cells that is fully compatible with the widely used 10x Genomics Chromium platform and standard Cell Ranger–based pipelines^16^. Our approach captures both short and long polyadenylated and non-polyadenylated RNAs using a minimal set of biochemical and computational modifications, enabling broad adoption without sacrificing throughput, accessibility, or interoperability with existing pipelines and datasets.

We applied this framework to over 500,000 single cells spanning diverse biological systems. In peripheral blood mononuclear cells (PBMCs), we detected robust cell-type–specific expression of non-coding RNAs, including miRNAs, tRNAs, and lncRNAs, and revealed the regulatory architecture of coding and non-coding co-expression modules. In cells infected with a non-polyadenylated RNA virus, we captured both host and viral transcripts, revealing antiviral programs otherwise missed by polyA-based protocols. In the developing human brain, we uncovered hundreds of dynamically regulated non-coding RNAs that trace developmental transitions and neuronal lineage specification.

Altogether, our work establishes a robust, accessible, and scalable strategy for total RNA profiling at single-cell resolution. Incorporating non-coding RNAs into single-cell datasets provides a foundation for deeper biological insights, improved computational modeling, and enhanced understanding of cell states across diverse biological contexts.

## Results

### TotalX streamlines high-throughput detection of non-coding RNA in single cells

To enable the robust detection of non-coding RNA species in a scalable single-cell framework, we adapted the principles of Smart-seq-total to a droplet-based 10x Genomics 3′ chemistry, resulting in a method we term TotalX (Smart-seq-total in 10X Chromium) (**Fig. 1a**). TotalX utilizes a custom template-switching oligo (dUTSO) and uracil-DNA glycosylase (UDG) digestion post–reverse transcription, as established in Smart-seq-total^10^. It further integrates Cas9-mediated ribosomal RNA (rRNA) depletion at the pre-amplified cDNA stage (DASH) (**Extended Fig. 1a-c**), while preserving full compatibility with standard 10x Genomics hardware and software (**Extended Fig. 1d-e and Supplementary Information**). In addition, we use custom oligos mimicking 10x Genomics barcode structure to index small fragments through direct amplification (**Supplementary Table 1**, see Methods).

**Figure 1.**
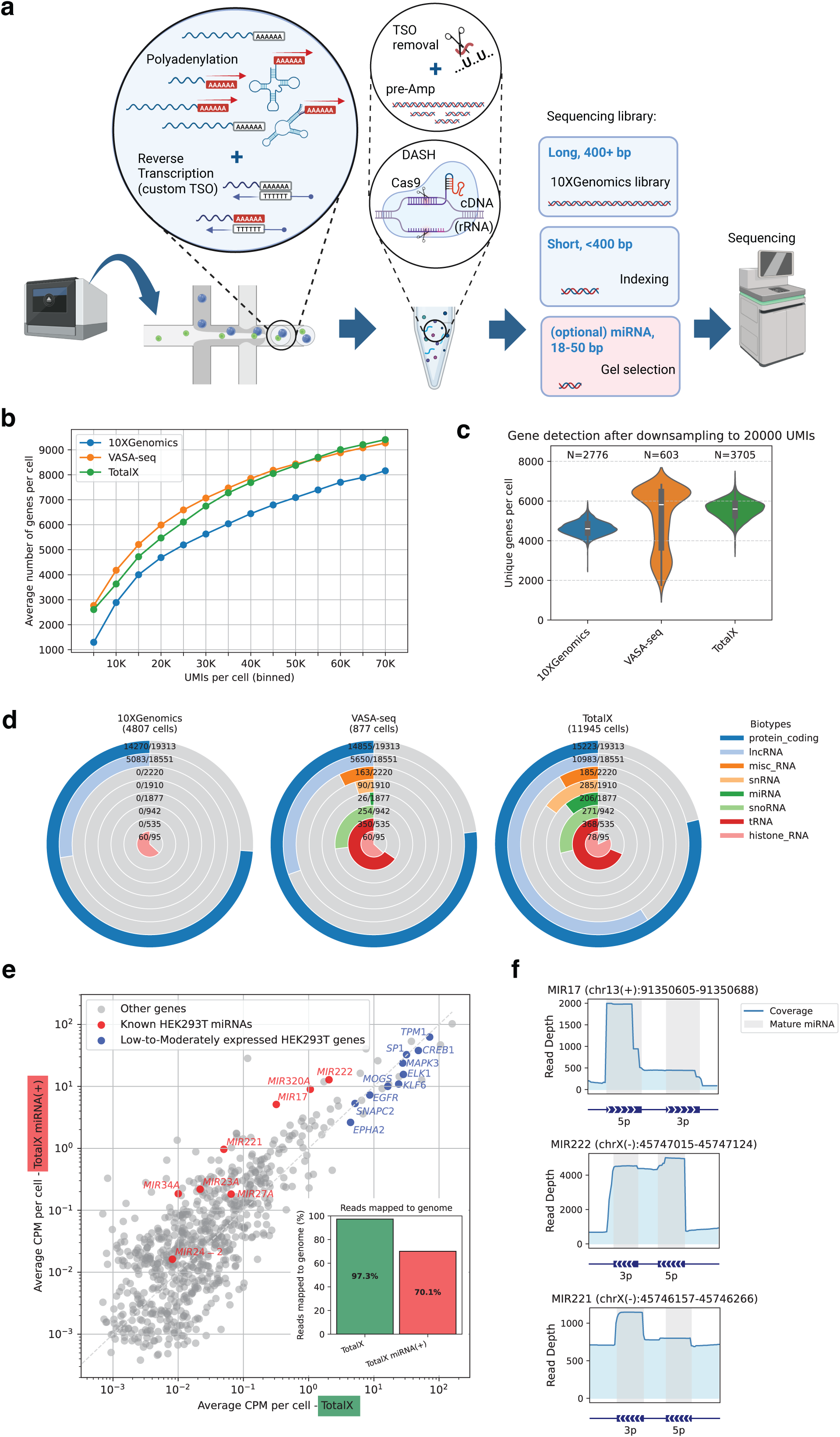
TotalX enables scalable detection of coding and non-coding RNAs, including miRNAs, in single cells. **a. Schematic overview of the TotalX protocol.** Total RNA is polyadenylated and reverse-transcribed using a custom template-switching oligo (dUTSO). After reverse transcription, the TSO is digested with UDG and rRNA is depleted at the pre-amplified cDNA level using Cas9-based DASH. Long (>400 bp) and short (<400 bp) fragments are indexed separately, with optional inclusion of a gel-purified miRNA fraction (~18–50 bp) and pooled for sequencing. The schematic was designed using https://BioRender.com. **b. Gene detection efficiency across technologies.** Comparison of average number of genes per cell as a function of UMIs for TotalX (green), VASA-seq (orange), and 10xGenomics 3′ chemistry (blue), across binned depths. **c. Unique genes detected per cell after UMI downsampling**. Gene detection following normalization to 20,000 UMIs per cell. TotalX yields high gene complexity similar to VASA-seq and higher than standard 10xGenomics Chromium 3′ workflow. **d. Total number of unique genes detected per RNA biotype.** Radial plots show numbers of unique genes detected in a representative experiment for each method, broken down by RNA biotype: protein-coding, lncRNA, miscRNA, miRNA, snoRNA, snRNA, tRNA, and histone RNA. Ratios represent the proportion of detected genes relative to the total number of annotated genes within each biotype. Only genes detected in 10 or more cells were counted. **e. Improved detection of miRNAs using mixed library input.** Scatter plot shows average CPM per cell of TotalX alone (x-axis) vs. TotalX with added miRNA fraction [TotalX-miRNA(+)] (y-axis) in HEK293T cells. Known HEK293T-specific miRNAs (red) reach expression levels comparable to low/moderate protein-coding genes (blue). Inset: proportion of reads mapping to genome, indicating a tradeoff with miRNA inclusion. **f. Coverage profiles for selected miRNAs.** Read depth plots for *MIR17*, *MIR222*, and *MIR221* showing mature miRNA arms (grey regions).

We benchmarked TotalX against VASA-seq, a high-performing non-coding RNA method^11^, and the standard 10x Genomics 3′ platform. TotalX achieves a comparable number of genes per UMI to VASA-seq, with a similar slope of gene recovery as a function of sequencing depth (**Fig. 1b**). After normalizing to 20,000 UMIs per cell, TotalX detects a comparable number of genes per cell to VASA-seq, while preserving the throughput of 10x Genomics Chromium droplet system with over 11,000 cells profiled in one experiment (**Fig. 1c and Extended Fig. 2a-b**). This enabled TotalX to capture a broader diversity of non-coding RNAs—including lncRNAs, miRNAs, snoRNAs, snRNAs, tRNAs, and histone RNAs—and to yield a higher number of unique genes per biotype within a single experiment (**Fig. 1d**).

**Figure 2.**
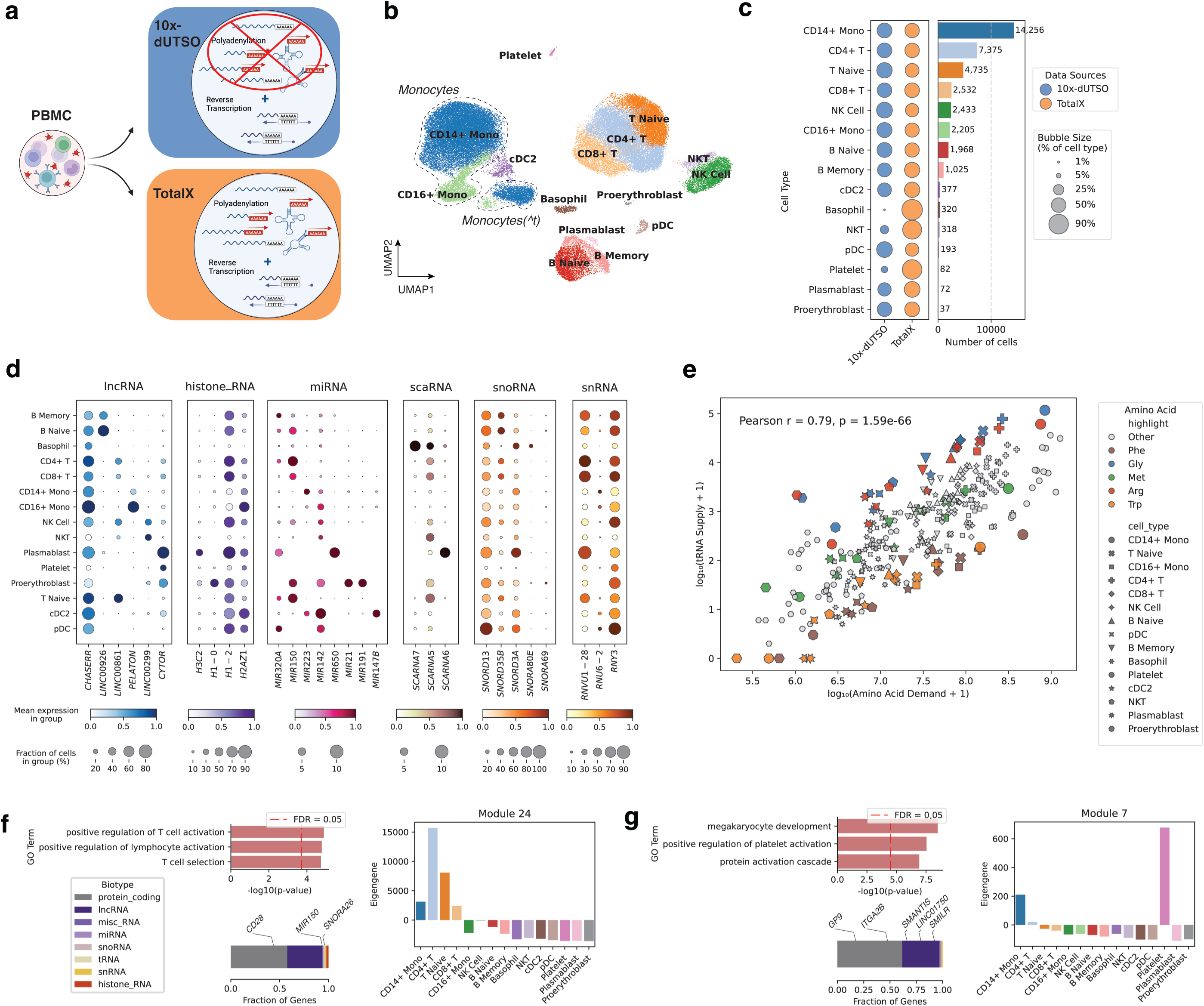
TotalX enables cell-type–specific profiling of non-coding RNAs in human PBMCs. **a. Schematic comparison of protocols applied to PBMCs.** TotalX incorporates enzymatic polyadenylation followed by reverse transcription using a custom TSO (10x-dUTSO). A non-polyadenylated 10x-dUTSO protocol is used for comparison. The schematic was designed using https://BioRender.com. **b. UMAP projection of TotalX-profiled PBMCs.** Cell types were identified using protein-coding gene expression and canonical markers. Two monocyte states are shown: regular and a transitional state (^t), each outlined by dashed-line contours. **c. Cell type frequencies across protocols**. Bar plot and bubble chart compare the number and relative proportion of recovered cell types between TotalX and 10x-dUTSO datasets. Bubble size reflects the fraction of each cell type within each dataset. Bar plot shows the number of cell detected in both datasets. **d. Cell-type–specific expression of non-coding RNAs**. Dot plots show representative lncRNA, histone RNA, miRNA, scaRNA, and snoRNA markers across annotated immune cell types. **e. Relationship between tRNA supply and amino acid demand**. Scatterplot shows global correlation across cell types (Pearson *r* = 0.79, *p* = 1.59e–66). Individual amino acids are highlighted; Arginine and Glycine show more supply, while Tryptophan and Phenylalanine show undersupply compared to other amino acids. **f. T cell–enriched gene co-expression module (Module 24)**. Top: GO enrichment terms for protein-coding genes in the module. Bottom left: Biotype composition of the module; Bottom right: module expression across cell types. Module includes *MIR150, SNORA26*, and *CD28*. **g. Platelet-specific gene co-expression module (Module 7).** Top: GO enrichment reveals association with platelet activation and megakaryocyte development. The module is composed largely of lncRNAs (e.g., *SMANTIS, SMILR, LINC01750*) co-expressed with *GP9* and *ITGA2B*.

To deepen detection of short RNA fragments, we performed size-selection for miRNA-enriched cDNA (~18–50 bp) and mixed this with longer fragments during library prep (**Extended Fig. 1b–c**). All RNAs are tagged using 10x Genomics-based UMI incorporation and thus the final libraries, even enriched, are demultiplex to count unique RNA molecules. This optional “miRNA(+)” strategy significantly increased detection of endogenous HEK293T-enriched miRNAs^17^ (**Fig. 1e-f**). The miRNAs were detected at the expression levels comparable to low-to-moderately expressed protein-coding genes, indicating strong capture sensitivity. We observed a tradeoff, however: the inclusion of size-selected short RNA fragments led to ~30% decrease of reads confidently mapped to the genome and transcriptome (**Fig. 1e**), as shorter reads are more likely to map ambiguously or represent adaptors with no insert (**Extended Fig. 2c-e**). Nevertheless, TotalX-miRNA(+) still retained strong detection of both long and short non-coding RNA, outperforming TotalX baseline version in biotype diversity (**Extended Fig. 2f–g**).

Read coverage profiles for individual miRNA genes, such as *MIR17*, *MIR221*, and *MIR222*, revealed precise detection of mature miRNA arms (5p or 3p) by TotalX (**Fig. 1f**). These profiles confirm that TotalX not only captures miRNA presence, but also preserves their biologically relevant processing patterns, which could be informative for downstream functional analysis and interpretation.

### Cell-type–resolved profiling of non-coding RNAs in human PBMCs

We next applied TotalX to human peripheral blood mononuclear cells (PBMCs) to evaluate its ability to profile coding and non-coding RNAs across diverse immune cell types (**Fig. 2a**). Using protein-coding genes for initial clustering and annotation (**Extended Fig. 3a**, Methods), we identified major expected populations including T and B lymphocytes, monocytes, NK cells, dendritic cells, plasmablasts, and proerythroblasts.

**Figure 3.**
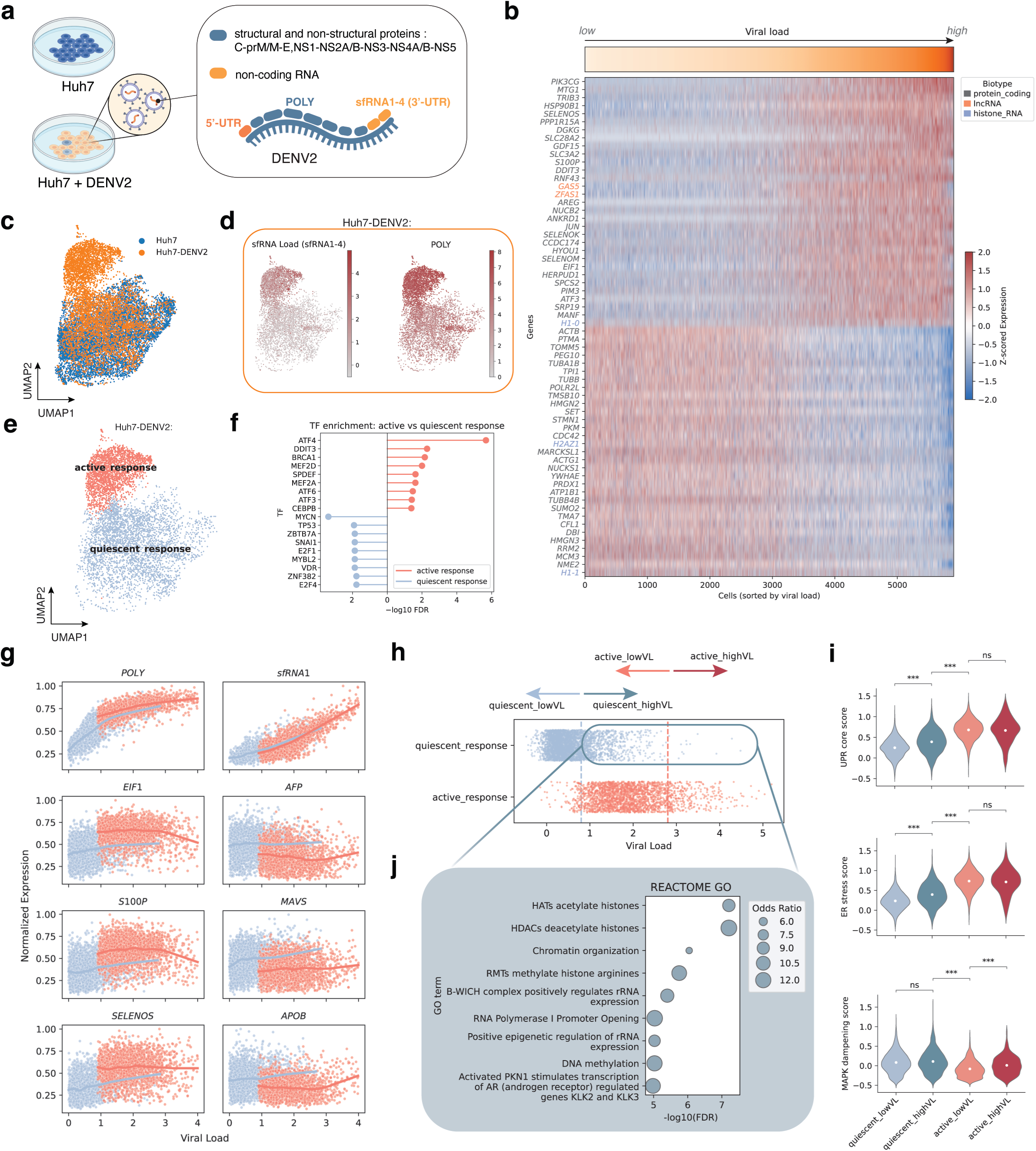
Distinct infection states and host responses in DENV2-infected cells. **a. Overview of DENV2 genome structure and infection design**. The positive-strand RNA genome encodes a polyprotein comprising structural (C-prM/M-E) and non-structural proteins (NS1–NS5), flanked by 5′ and 3′ UTRs. The 3′UTR gives rise to four highly structured, non-coding subgenomic RNAs (sfRNA1–4). Huh7 cells were infected with DENV2 and profiled using TotalX. **b. Host gene expression correlation with viral load.** Heatmap of coding and non-coding host genes (rows) sorted by increasing viral load across infected cells (columns). *POLY* and *sfRNA* levels contribute to a composite viral load score. **c. UMAP projection of DENV2-infected and uninfected cells.** Clustering based on host protein-coding gene expression. **d. Abundance of *sfRNA1* and *POLY* in infected cells**. Feature plots confirm detection of viral non-coding and coding transcripts at the single-cell level. **e. Infection states resolved by unsupervised clustering of single-cell host transcriptomes.** Infected cells separate into two transcriptionally distinct clusters: active and quiescent host response. **f. Transcription factor (TF) enrichment analysis.** Gene set enrichment analysis (GSEA) shows activation of ATF4 and DDIT3, among other, in infected cells with active host response. **g. Examples of gene expression vs. viral load.** *POLY* and *sfRNA1* increase monotonically with load, while host stress response and interferon genes diverge between active and quiescent states. Normalized expression values are scaled between 0 and 1 for each gene. Dots represent single cells; lines indicate LOWESS-smoothed trends for each cluster. Colors denote distinct clusters as indicated in the legend. **h. Viral load distribution across states.** Quiescent and active infection groups both contain cells with low and high viral load, indicating that viral load alone does not determine host response. **i. Enrichment of chromatin and RNA regulation pathways**. Reactome GO enrichment showing histone deacetylation and acetylation signatures in the quiescent response, high viral load (quiescent_highVL) state. **j. Module scores for infection-associated programs.** Violin plots show activation of unfolded protein response (UPR), ER stress, granule formation, and MAPK signaling across groups. UPR/ER stress increases in quiescent cells with high viral load (quiescent_highVL) cells, but MAPK signaling remains suppressed.

Compared to a version of the same protocol lacking the polyadenylation step (10x-dUTSO), TotalX recovered a similar diversity of immune cell types (**Fig. 2b**) and allowed for more consistent detection of low RNA content cell types, including platelets and basophils, though we do not claim comprehensive detection of rare populations (**Fig. 2c**). Importantly, we observed that monocytes were sensitive to protocol conditions, especially the presence of ATP used during the enzymatic polyadenylation step. In agreement with prior reports^18^, the presence of ATP led to a transcriptional transition state in monocytes characterized by downregulation of *LYZ* and upregulation of *JARID2* and *GAB2*, accompanied by a global reduction in gene expression and a partial loss of canonical identity (**Extended Fig. 3b–g**). Other cell types were not affected by the polyadenylation conditions in our assay.

TotalX enabled detection of non-coding RNAs that are consistently expressed in defined immune cell populations, supporting their role in maintaining cell identity. Wilcoxon rank-sum testing identified hundreds of differentially expressed non-coding RNAs (log2(FC)>1, adjusted p-value < 0.05) across cell types (**Extended Fig. 4a–b, Supplementary Table 2**). For example, *MIR650* was enriched in plasmablasts, *MIR147B* in cDC2s, *MIR150* in T cells, proerythroblasts, and naïve B cells. Several lncRNAs also showed cell-type enrichment, including *LINC00299* in NK and NKT cells, and *PELATON* in monocytes. We additionally detected structured non-coding RNAs such as *SCARNA6* (plasmablasts) and *SNORD13* (pDCs) (**Fig. 2d**), suggesting broad recovery of functionally relevant small RNAs.

**Figure 4.**
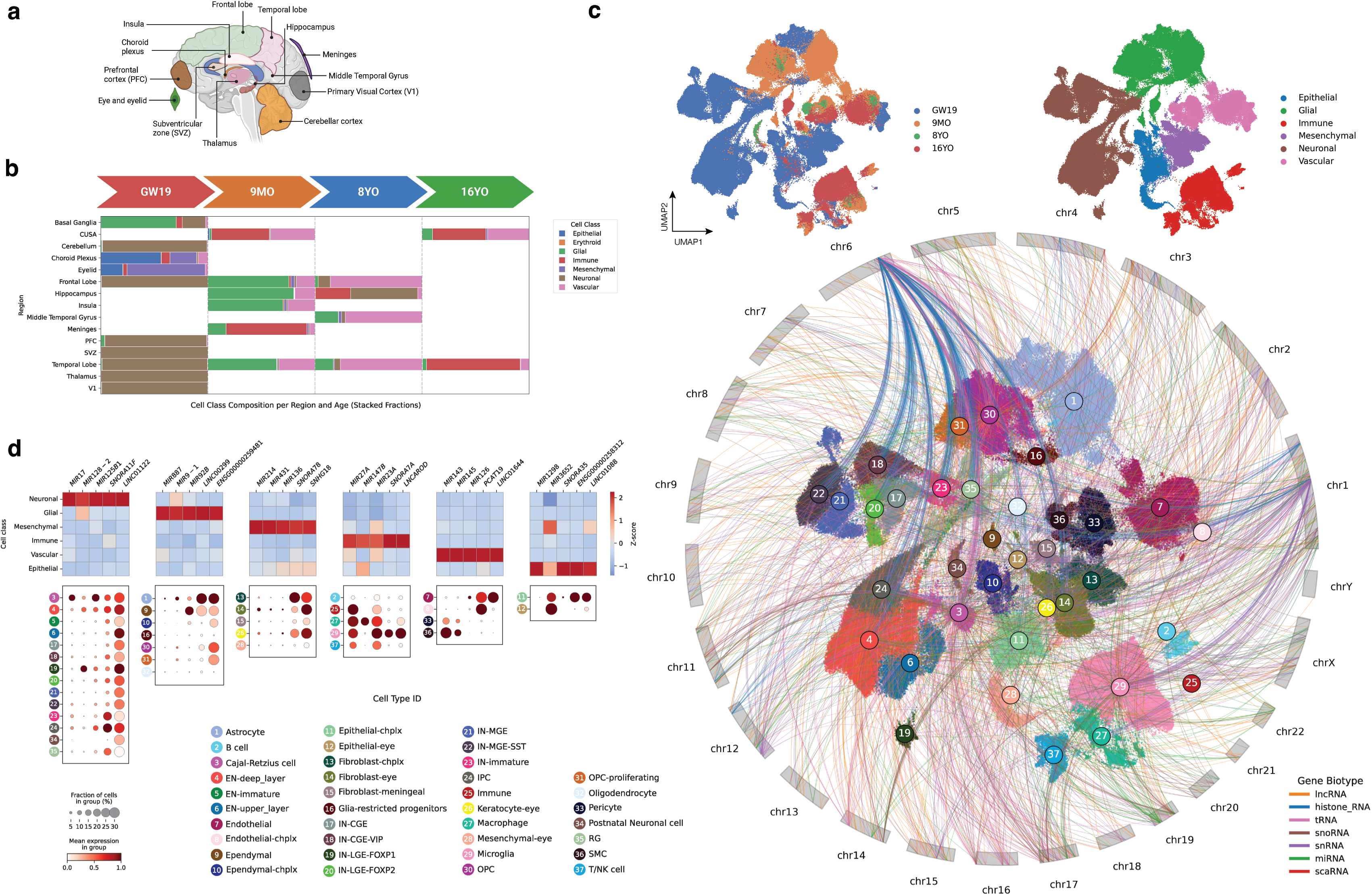
TotalX reveals cell-type–specific expression of non-coding RNAs across diverse regions of the developing human brain. **a. Schematic of sampled brain regions.** Dissected regions include cortex (frontal, temporal, and prefrontal), subventricular zone (SVZ), hippocampus, thalamus, cerebellar cortex, choroid plexus, insula, meninges, middle temporal gyrus, and eye tissue. Developmental timepoints span gestational week 19 (GW19), 9 months postnatal, 8 years, and 16 years. **b. Cell class composition across age and region.** Stacked bar plot showing the distribution of major cell classes (e.g., neuronal, glial, epithelial, immune) across sampled regions at each developmental stage. **c. UMAP projection of 301,515 single cells profiled using TotalX.** Cells are colored by cell type (main plot), cell class (top right), and age (top left). Arcs connect the most cell-type– specific non-coding RNA markers (top 400 per cell type, selected based on log fold change > 1 and adjusted p-value < 0.01)—including miRNAs, lncRNAs, snRNAs, and histone RNAs—to their genomic coordinates. Arcs are colored by gene biotype, illustrating relationships among RNA biotypes, chromosomal loci, and cell classes. **d. Selected cell-type–specific non-coding RNA markers.** Dot plots display expression of non-coding RNAs across brain cell types, grouped by broad cell classes.

Because TotalX captures tRNAs, we next asked whether there is alignment between tRNA availability and amino acid demand based on codon usage (Methods). Across cell types, we observed a strong correlation (Pearson r = 0.79, p = 1.59e–66). However, individual amino acids such as arginine and glycine exhibited relatively higher tRNA supply compared to others, while tryptophan and phenylalanine were more limited (**Fig. 2e**, **Extended Fig. 4c**). These modest deviations indicate that while codon usage strongly predicts tRNA availability, there may be additional influences shaping tRNA pools.

To investigate how non-coding RNAs are co-regulated with protein-coding genes, we performed weighted gene co-expression network analysis (WGCNA, see Methods) across all TotalX-profiled PBMCs. We identified over 30 gene modules comprising both coding and non-coding transcripts (**Extended Fig. 4d–e, Supplementary Table 3**). For example, Module 24, enriched in T cells and CD14+ monocytes, included *MIR150*, *SNORA26*, and *CD28*, and was associated with GO terms related to lymphocyte activation (**Fig. 2f**). Module 7, a platelet-specific module, included lncRNAs such as *SMANTIS*, *SMILR*, and *LINC01750*, co-expressed with canonical platelet transcripts *GP9* and *ITGA2B*, and enriched for terms related to platelet activation and megakaryocyte development (**Fig. 2g**).

These results highlight that non-coding RNAs are not only detectable but systematically co-expressed with functional coding genes in a cell-type–specific manner, suggesting their potential roles in immune cell identity and function.

### Co-detection of non-polyadenylated viral transcripts and host transcriptome in DENV2-infected cells

We next evaluated whether TotalX enables simultaneous detection of non-polyadenylated viral RNAs and host transcripts within individual cells. As a test case, we profiled human Huh7 cells infected with dengue virus serotype 2 (DENV2)—a non-polyadenylated flavivirus whose RNA genome encodes a single polyprotein composed of both structural/non-structural proteins (C-prM/M-E-NS1-NS5) and a family of structured, non-coding RNAs (sfRNA1–4) that originate from the 3′ untranslated region (UTR)^19,20^ (**Fig. 3a**, **Extended Fig. 5a**).

**Figure 5.**
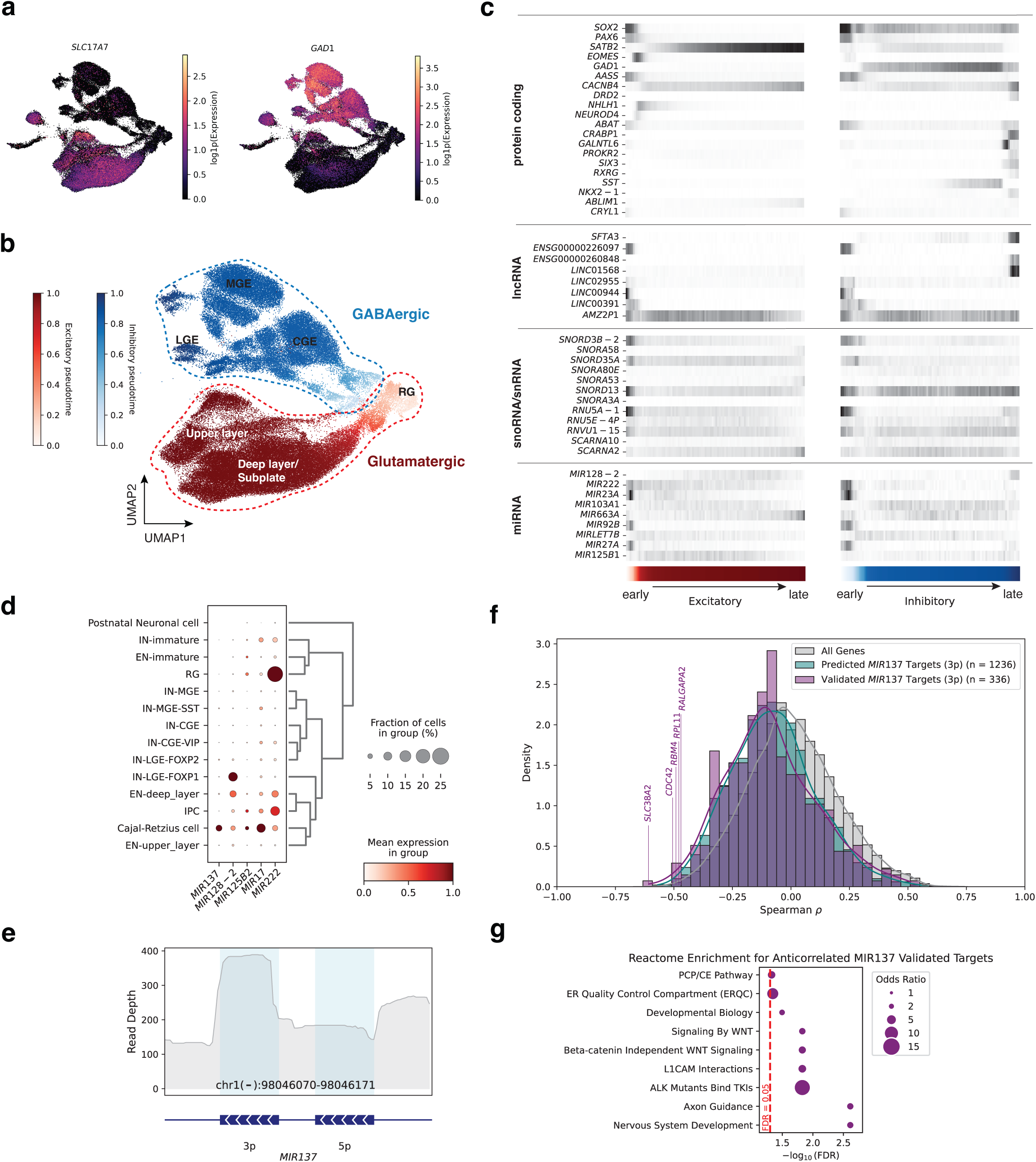
Cell-type–resolved expression of disease-relevant miRNAs across developing neuronal lineages. **a. UMAP of neuronal lineages annotated by marker gene expression.** Expression of canonical glutamatergic (*SLC17A7*) and GABAergic (*GAD1*) markers delineates excitatory and inhibitory trajectories across developing brain regions, including cortical, subplate, and ganglionic eminence–derived neuroblasts. **b. Pseudotime trajectories of glutamatergic and GABAergic neuronal lineages.** Pseudotime was inferred using PAGA. **c. Selected gene expression across neuronal progenitors and mature neuron types**. Selected gene expression across neuronal progenitors and mature neuron types. The heatmap shows relative expression of protein-coding genes, lncRNAs, snoRNAs, snRNAs, and miRNAs across major excitatory and inhibitory neuronal lineages. Expression values represent normalized means across 100 pseudotime-defined bins. The color bar indicates pseudotime, consistent with panel b. **d. Dot plot of disease-associated miRNAs abundance across neuronal cell types.** **e. Read coverage profile of *MIR137* gene locus.** Aggregate read depth across the *MIR137* gene region (chr1(-):98,046,070–98,046,171), showing precise coverage of mature 3p arm by TotalX in developing neuronal lineages. **f. Spearman correlation between *MIR137* and its targets.** Histogram of Spearman correlation coefficients across *MIR137* and all other genes. Some *MIR137* validated targets are significantly anticorrelated (adjusted p-value<0.05), consistent with post-transcriptional repression. Vertical lines and gene names denote top anticorrelated protein coding genes. **g. GO enrichment for *MIR137* targets**. Reactome GO analysis of negatively correlated (Spearman rho < −0.3. adjusted p-value < 0.01) validated *MIR137* targets reveals enrichment for pathways related to nervous system development, WNT signaling, and axon guidance.

We applied TotalX to infected and mock-treated Huh7 cells and successfully recovered viral transcripts including both protein-coding regions (*POLY*) and structured non-coding RNAs (*sfRNAs*) in the infected cells. The number of viral UMIs per 10,000 host UMIs exceeded prior reports^21^, highlighting TotalX’s sensitivity to short, structured RNAs (**Extended Fig. 5b**). To quantify infection per cell, we defined a viral load score based on summed expression of *POLY* and *sfRNA1–4*. This score correlated with broad host transcriptomic changes across both protein-coding and non-coding genes (**Fig. 3b**).

Notably, infected cells partitioned into two transcriptional states—“active response” and “quiescent response”—based on host gene expression (**Fig. 3c–e**). Despite similar viral levels, cells in the quiescent response state failed to mount the canonical antiviral response and transcriptionally looked similar to uninfected cells (**Fig. 3c–e**). This heterogeneity is consistent with the infection conditions used (MOI = 1, 48 h post-infection; Methods), which are expected to yield a mixture of uninfected cells, secondary-infected cells, and more advanced primary-infected cells within the population. Transcription factor (TF) enrichment analysis using GSEA^22^, showed that active response was associated with ATF4 and DDIT3 activation— regulators of the unfolded protein response (UPR) and ER stress (**Fig. 3f**) —consistent with prior studies^23^. In contrast, quiescent cells did not show upregulation of these targets.

Surprisingly, we found a subset of “quiescent highVL” cells—those with high viral load but suppressed host activation. While these cells contained abundant levels of both *POLY* and *sfRNA* transcripts (**Fig. 3g–h**), they maintained baseline (uninfected) levels of *EIF1* and *S100P* and also showed higher levels of *AFP, APOB* and *MAVS* compared to the “active response” group (**Fig. 3g**). In addition, the “quiescent highVL” cells exhibited muted inflammatory and stress-response signatures, including downregulation of MAPK signaling (**Fig. 3i**). This observation prompted us to explore possible mechanisms underlying the suppression of host responses in these cells. Although prior studies have shown that *sfRNA* can inhibit host antiviral defenses^24,25^, the precise factors at play in our system remain unclear. To investigate further, we performed differential expression analysis across high (highVL) and low viral load (lowVL) cells within both active and quiescent cell response states (**Extended Fig. 6d, Supplementary Table 4**). In the “quiescent highVL” group, we observed strong enrichment of chromatin regulatory signatures, including histone deacetylase (HDAC) and histone acetyltransferase (HAT) pathways (**Fig. 3j**), implicating epigenetic remodeling in the suppression of host responses. Previous studies have shown that HDAC inhibition during DENV2 infection can modulate inflammatory cytokine production^26^; here, we provide a transcriptome-wide, single-cell–level quantification of this effect, highlighting a potential mechanism of immune evasion through chromatin-based silencing.

### Region- and cell-type–specific expression of non-coding RNAs in the developing human brain

To chart the cellular and spatial distribution of non-coding RNAs in the human brain, we applied TotalX to neurosurgical and fetal tissue collected across four developmental stages: gestational week 19 (GW19), 9 months (9MO), 8 years (8YO), and 16 years (16YO) (**Fig. 4a**). The dataset includes 22 biological samples from 4 individuals, covering a broad array of brain regions, including the prefrontal (PFC) and primary visual (V1) cortices, thalamus, hippocampus, cerebellum, insula, meninges, choroid plexus, ventricular zones, and ocular tissue (**Fig. 4b and Extended Fig. 7a-b**).

We profiled 301,515 single cells using the TotalX-miRNA(+) protocol, which enabled robust detection of diverse non-coding RNA biotypes—including miRNAs, lncRNAs, snoRNAs, snRNAs, tRNAs, and histone mRNAs—across a broad range of cell types and developmental stages. Dimensionality reduction revealed distinct transcriptional clusters corresponding to canonical neural (including inhibitory [IN], and excitatory [EN] neurons), glial, immune, and non-neural cell types, with clear separation by developmental stage and brain region (**Fig. 4c**). Cell types were annotated based on established coding gene markers (**Extended Fig. 7c**)^27–30^ and further characterized by their non-coding RNA expression profiles.

Non-coding RNA expression varied markedly across cell types and developmental stages. Histone mRNAs were highly expressed in rapidly dividing progenitors such as radial glia, while snRNAs and lncRNAs were enriched in immune and differentiated neuronal populations. In total, we identified hundreds of differentially expressed non-polyadenylated transcripts, revealing their widespread contribution to cell identity and developmental state (**Extended Fig. 7d)**.

To examine how non-coding RNA expression relates to genome architecture, we mapped the top cell-type–specific non-coding RNA markers to their chromosomal coordinates (**Fig. 4c**). This analysis uncovered biotype- and lineage-specific enrichment patterns: for instance, a histone gene cluster on chromosome 6 was predominantly expressed in progenitors, while a snRNA cluster on chromosome 1 was enriched in microglia. These patterns point to coordinated regulation of non-coding RNAs across both genetic context and cell type.

We next looked for cell-type–specific non-coding RNA markers. **Fig. 4d** highlights selected non-coding RNAs across major cell types and RNA biotypes. Dot plots and heatmaps show the expression of representative lncRNAs, miRNAs, and snoRNAs across neuronal, glial, immune, epithelial, and mesenchymal populations. Several miRNAs exhibited striking cell-type specificity. For example, *MIR17* was highly expressed in Cajal-Retzius cells^31,32^ and deep-layer excitatory neurons (EN-deep_layer), while *MIR128-2* was detected in the same populations but showed peak expression in lateral ganglionic eminence-derived interneurons (IN-LGE-FOXP1). *MIR9-1* was enriched in astrocytes, *MIR92B* in ependymal cells, and *MIR125B1* in early neurons. We also observed *MIR147B* expression in macrophages but not microglia, and broad expression of *MIR23A* and *MIR27A* across immune, stromal, and epithelial populations, and were also detected in a subset of radial glia (**Extended Fig. 7e**).

Beyond miRNAs, other biotypes displayed similarly specific expression patterns. For example, *LINC00299* was enriched in astrocytes and ependymal cells, *SNORA7A* and *LNCAROD* in microglia, and *SNORA11F* in neurons. *ENSG00000258312* marked choroid plexus ependymal cells, while *LINC01644* and *PCAT19* were detected in endothelial cells. *SNHG18* and *SNORA78* were broadly expressed across mesenchymal and epithelial progenitor populations (**Fig. 4d**).

Together, these data show that non-coding RNA, along with coding genes, exhibit distinct and dynamic patterns of cell-type and temporal specificity. Their abundance deepens the layered regulatory complexity of brain development and highlights the utility of total RNA profiling for resolving the non-coding transcriptome at single-cell resolution.

### Non-coding RNA dynamics across developing neuronal and glial lineages

To investigate the role of non-coding RNAs in neuronal lineage commitment, we focused on developing excitatory and inhibitory neurons (**Fig. 5a–b**). Using partition-based graph abstraction (PAGA) and pseudotime alignment, we reconstructed developmental trajectories from radial glia (RG) toward both glutamatergic neuronal lineages, and from ganglionic eminence progenitors toward GABAergic neuronal lineages^33^. While most neurons in our dataset remain in immature states, we observed robust expression of canonical early differentiation markers. These included *EOMES* in intermediate progenitors, *DCX* in newborn neurons, and *SATB2* and *SLC17A7* in developing excitatory neurons, as well as *DLX1*, *DLX2*, and *GAD1* in GABAergic interneuron precursors (**Fig. 5a–b**, **Extended Fig. 7b** and **8a–b**), consistent with previous studies of the developing human brain^27,34^.

Along these trajectories, we identified dozens of developmentally dynamic non-coding RNAs— including miRNAs, lncRNAs, and snRNAs (**Fig. 5c**). Several small RNAs with known neurological relevance, such as *MIR222*, *MIR103A1*, and *MIR128-2*^35–37^, exhibited transient expression peaks in early neurons and declined with maturation, suggesting potential roles in fate specification and lineage progression.

We performed similar analyses for glial trajectories, reconstructing differentiation paths from RG into glia-restricted progenitor cells^38^ to astrocytes or oligodendrocyte precursor cells (OPCs), and mature oligodendrocytes (**Extended Fig. 8c–f**). We observed comparable dynamic patterns of non-coding RNA expression along glial lineages. *LINC00299* was upregulated in late-stage astrocytes, while *MIR219A2HG* showed increased expression at later stages of OPC differentiation. *MIR143* was enriched in mature astrocytes, whereas *MIR568* displayed a dynamic trajectory—highly expressed in radial glia, downregulated in astrocyte progenitors, and then re-expressed during astrocyte and OPC maturation. Notably, *MIR23A* was also expressed in RG and re-emerged in mature oligodendrocytes, suggesting a possible reuse of regulatory programs at distinct stages of glial development.

### miRNA dynamics and target repression across developing neuronal trajectories

Cajal-Retzius (CR) cells are one of the earliest-born neurons in the developing cortex, though they largely disappear by birth. During development, they occupy the outermost cortical layer I, and guide radial neuron migration by secretion of reelin (RELN)^39,40^. Among the developing neuronal populations, CR cells emerged as a hotspot of miRNA expression, displaying higher overall miRNA abundance compared to other neuronal subtypes (**Extended Fig. 9a**). This suggests that CR cells—though transient—may rely on enhanced post-transcriptional regulation during early cortical development. Within this population, *MIR137* stood out for its specificity: it was highly expressed in CR cells and nearly absent from other neuronal subtypes (**Fig. 5d**). *MIR137* is also one of the strongest GWAS-associated loci for schizophrenia and intellectual disability^41,42^, with established roles in synaptic development and epigenetic regulation^43^. Additional miRNAs, including *MIR17* and *MIR222*—previously linked to neuronal survival and axon outgrowth^44,45^—also showed enriched expression across several populations during early neurogenesis.

To investigate whether these miRNAs exert post-transcriptional control within the same cells, we examined the expression of their validated targets. For *MIR137*, TotalX read coverage revealed detection of both 5p and 3p arms, with expression dominated by the 3p arm (**Fig. 5e**)—consistent with prior findings showing that *miR-137-3p* accounts for 97% of all *miR-137*-specific reads across 71 sequencing experiments compiled by miRBase^46^. *MIR137* was selectively expressed in early neurons, particularly CR cells, whereas many of its known targets—genes involved in cell cycle progression, chromatin remodeling, RNA splicing, and synaptic vesicle transport—were enriched in neuronal progenitors and displayed mutually exclusive expression with *MIR137*. Spearman correlation analysis confirmed a significant anticorrelation between *MIR137* and some of its validated targets (adjusted p-value<0.05) (**Fig. 5f**), consistent with miRNA-mediated repression^47^. Among the top anticorrelated genes were *SLC38A2* and *CDC42*, the latter of which has previously been shown to be downregulated by *MIR137* in cancer cells^48^. GO enrichment analysis further implicated *MIR137* targets in nervous system development and Wnt signaling (**Fig. 5g**), underscoring their relevance to early neurogenesis.

We performed similar analyses for *MIR17*, *MIR125B-2*, and *MIR128-2* (**Extended Fig. 9c-e**), and found that both validated and predicted targets^47,49^ were consistently anticorrelated with miRNA expression (adjusted p-value<0.05). These patterns suggest widespread post-transcriptional repression of developmental gene programs by miRNAs during early neuronal subtype differentiation.

Together, these findings support a critical role for non-coding RNAs—particularly miRNAs—in shaping early neuronal lineage commitment, and position CR cells as a unique miRNA-regulated population during cortical development. More broadly, they highlight the power of TotalX to resolve dynamic, biotype-diverse regulatory programs at single-cell resolution and to uncover functional miRNA–target relationships.

## Discussion

This study introduces TotalX, a scalable and accessible framework for total RNA profiling in single cells that enables the simultaneous detection of both polyadenylated and non-polyadenylated transcripts. By building upon the widely adopted 10x Genomics Chromium platform with minimal protocol and software modifications, TotalX overcomes long-standing limitations in single-cell transcriptomics where non-coding RNAs—particularly short, structured, or non-polyadenylated species—have remained largely inaccessible. Through benchmarking, optimization, and application across diverse biological contexts, we demonstrate that TotalX provides a biotype-rich molecular view of individual cells while preserving throughput and compatibility with existing pipelines.

Using TotalX, we detected diverse classes of non-coding RNAs, including miRNAs, tRNAs, lncRNAs, snoRNAs, snRNAs, histone RNAs, at single-cell resolution across immune cells, virally infected hepatocytes, and the developing human brain. In PBMCs, we identified non-coding RNA markers of cell identity and co-expression modules linking coding and non-coding genes. In DENV2-infected cells, TotalX captured both coding and structured viral RNAs and identified a transcriptionally quiescent but high viral load infection state potentially linked to chromatin regulation. In the developing brain, we mapped the landscape of non-coding RNA expression across regions and developmental timepoints, uncovering biotype-specific enrichment patterns and chromosomal localization of cell-type–restricted non-coding RNAs. These findings highlight the biological value of simultaneously capturing coding and non-coding RNA programs in single cells.

CR cells, among the earliest-born neurons in the cerebral cortex, play a foundational role in brain development by secreting reelin and guiding the laminar architecture^39^. Although transient, they scaffold the future cortex. Our finding that CR cells exhibit enriched expression of *MIR137*—a miRNA strongly associated with schizophrenia risk^41^—raises the possibility that post-transcriptional dysregulation in this population could have lasting consequences for cortical circuit assembly. Because *MIR137* regulates genes involved in chromatin remodeling, synaptic function, and neuronal maturation, its misexpression during the narrow developmental window when CR cells are active may contribute to neurodevelopmental disorders. These observations potentially position CR cells as both structural architects and a regulatory vulnerability point in psychiatric disease.

A key advantage of TotalX lies in its ability to resolve dynamic expression of small RNAs— such as miRNAs—and link them to functional outcomes. For instance, in developing neuronal lineages, we identified temporally restricted expression of several miRNAs and demonstrated strong anticorrelation with its validated targets, suggesting direct post-transcriptional repression within the same cell. This level of resolution is difficult to achieve using traditional polyA-capture methods or indirect inferences from bulk data. More broadly, the ability to recover miRNA–target relationships, histone mRNA bursts, and coordinated tRNA-codon programs opens new avenues for studying cell state transitions, stress responses, and translational control.

Despite these advances, TotalX has limitations. The addition of short RNA libraries, while improving small RNA detection, can reduce the overall proportion of mappable reads and requires careful optimization of library mixing ratios. Detection of circular RNAs and very low abundance transcripts remains limited, and future protocol iterations may benefit from dedicated capture strategies or enrichment steps. In particular, low-abundance non-coding RNAs—such as certain lncRNAs, snoRNAs, and microRNAs—are also potentially underrepresented or missed entirely in the current workflow. Furthermore, although our modified Cell Ranger pipeline enables short read processing, specialized quantification tools may improve detection sensitivity and assignment accuracy for certain non-coding classes.

Looking ahead, total RNA profiling with TotalX opens new opportunities for comprehensive single-cell atlases, unbiased perturbation screens, and AI-driven models of cell identity. By expanding the scope of what is measurable in single cells, TotalX allows researchers to interrogate regulatory landscapes with unprecedented depth—supporting discovery of unanticipated transcripts and mechanisms. Critically, this method generates rich, integrative data fully compatible with widely used workflows, such as single-cell atlases and Perturb-seq screens on the 10x Genomics Chromium platform^50,51^. Resolving coding and non-coding elements simultaneously within single cells provides a more complete molecular phenotype, essential for uncovering subtle regulatory interactions, rare cell states, and context-specific responses.

## Methods

### HEK293T cell isolation

HEK293T cells were cultured in complete DMEM high glucose medium (Thermo Fisher Scientific, 11965092) supplemented with 10% fetal bovine serum (Thermo Fisher Scientific, 16000044), 1mM sodium pyruvate (Thermo Fisher Scientific, 11360070) and 100 μg/mL Penicillin/Streptomycin (Thermo Fisher Scientific, 15070063). Cells were dissociated using 0.25% Trypsin-EDTA (Thermo Fisher Scientific, 25200056) for 2-4 min at 37 °C and collected for analysis.

### Peripheral blood mononuclear cell isolation

Peripheral blood mononuclear cells (PBMCs) were isolated from a leukoreduction system (LRS) chamber collected from a healthy 61-year-old adult male donor at Stanford Blood Center. The LRS chamber product was diluted 1:4 in phosphate-buffered saline (PBS) supplemented with 2% fetal bovine serum (FBS; Thermo Fisher Scientific, 16000044) and layered onto Ficoll-Paque Plus (GE Healthcare, 17-1440-02) for density gradient centrifugation at 400 × g for 30 min at room temperature, with brake off. The mononuclear cell layer was collected, washed twice in PBS + 2% FBS, and residual red blood cells were lysed using ACK lysis buffer (Thermo Fisher Scientific, A1049201) for 3 min at room temperature. Cells were then washed, counted, and resuspended at 1 × 10 cells/ml in CryoStor CS10 cryopreservation medium (Stemcell Technologies, 07930), aliquoted into cryovials, and frozen at −80 °C in a controlled-rate freezing container before long-term storage in liquid nitrogen.

Prior to single-cell capture, cryopreserved PBMCs were rapidly thawed at 37 °C, transferred to pre-warmed RPMI 1640 medium (Thermo Fisher Scientific, 11875093) supplemented with 10% FBS, and washed twice in PBS + 0.04% BSA (Thermo Fisher Scientific, AM2616). Viability and concentration were assessed using the Luna automated cell counter (Logos Biosystems) with Trypan Blue exclusion. Final input cell concentrations were adjusted to 750–1,000 cells/µl.

### Dengue virus infection of Huh7 cells

Huh7 human hepatoma cells (Apath LLC) were cultured in Dulbecco’s modified Eagle’s medium (DMEM; Thermo Fisher Scientific, 11965092) supplemented with 10% fetal bovine serum (FBS; Thermo Fisher Scientific, 16000044), 1 mM sodium pyruvate (Thermo Fisher Scientific, 11360070), and 100 U/ml penicillin-streptomycin (Thermo Fisher Scientific, 15140122) at 37 °C in a 5% CO humidified incubator. Cells were seeded in 6-well plates and infected at ~70% confluence with dengue virus serotype 2 (DENV2; strain 16681) at a multiplicity of infection (MOI) of 1 as described in Zanini et al.^21^.

For infection, virus-containing media was diluted in serum-free DMEM and added to cells for 1 h at 37 °C with gentle rocking every 15 min. After incubation, the inoculum was removed, and cells were washed once with phosphate-buffered saline (PBS; ThermoFisher Scientific, 10010023) before replacing with fresh complete medium. Cells were incubated for 48 h post-infection to allow for robust viral replication. Mock-infected controls were treated identically with virus-free media.

At the endpoint, cells from two replicate wells (approximately 1 × 10 cells per well) were harvested by trypsinization (0.25% Trypsin-EDTA; Thermo Fisher Scientific, 25200056), pelleted by centrifugation (300 × g, 5 min), and resuspended in PBS for downstream processing. Cell viability was assessed using the Luna automated cell counter (Logos Biosystems), and cells were diluted to 750–1,000 cells/µl prior to encapsulation for single-cell TotalX.

### Human brain tissue processing

**Fetal brain samples** (19 gestational weeks) were obtained from Advanced Bioscience Resources (Newark, CA) and shipped overnight in cold preservation solution. All procedures were approved by the Stanford IRB and Stanford SCRO Panel. Intact samples were dissected into anatomical regions by licensed neuropathologists, minced using sterile razor blades, and digested in HBSS (ThermoFisher, 24020117) containing 10 mg/mL Liberase (Roche, 5401119001) and 200 µg/mL DNase I (Worthington, LS002007) for 40 min at 37 °C with gentle agitation. The digestion was repeated once. Samples were then incubated in Accutase (Innovative Cell Technologies, AT104) supplemented with 200 µg/mL DNase I for 15 min at 37 °C. Red blood cells were removed using Histopaque-1077 (Sigma, 10771) by layering the suspension at a 2:1 ratio and centrifuging at 400 × g for 30 min at 25 °C (low acceleration, no brake). The buffy coat was collected and washed in HBSS containing 0.1% polyvinyl alcohol (PVA; Sigma, P8136).

**Neurosurgical brain tissue samples** (from 9-month-old, 8-year-old, and 16-year-old individuals) were obtained during epilepsy surgeries at Stanford Hospital with informed consent under a protocol approved by the Stanford Institutional Review Board. Tissue was transferred on ice immediately from the operating room and processed within 1 h of resection. Samples were manually minced and enzymatically dissociated using papain (Worthington Biochemical, LK003176) or Accutase (Innovative Cell Technologies, AT104) supplemented with DNase I (Worthington, LS002007), depending on sample integrity and downstream application. Cell suspensions were filtered through 40 µm strainers (Corning, 352340) and used directly for downstream assays.

Following dissociation, cortical samples exhibited high levels of myelin debris and CD45 vascular macrophages. These populations were depleted in two steps. First, magnetic bead-based negative selection was performed using anti-CD45 microbeads (Miltenyi Biotec, 130-045-801), followed by 30 min incubation at 4 °C and magnetic separation using MS columns (Miltenyi Biotec, 130-096-433). Second, residual myelin was depleted by incubation with Myelin Removal Beads II following the manufacturer protocol. Post-depletion, cell suspensions were counted using a Luna automated cell counter (Logos Biosystems) and viability was assessed using LIVE/DEAD viability/cytotoxicity reagent (Thermo Fisher Scientific, L3224). Final cell concentrations were adjusted to 750–1,000 cells per µl prior to encapsulation.

### Single-cell total RNA sequencing with TotalX

Single-cell total RNA sequencing was performed using TotalX, a modified 10x Genomics Chromium 3′ v3.1 workflow incorporating enzymatic strategies adapted from Smart-seq-total^10^. The protocol enables the simultaneous capture of polyadenylated and non-polyadenylated transcripts through a co-incubation approach integrating 3′ polyadenylation, 5′ capping, and reverse transcription (RT) in a single reaction.

RT master mix was prepared by supplementing the standard 10x RT reagents with the addition of following components: E. coli poly(A) polymerase (New England Biolabs, M0276S; final 1 U/µl), ATP (10 mM), Vaccinia capping enzyme (NEB, M2080S; 0.1 U/µl), S-adenosylmethionine (SAM, 2 mM; NEB, M2080S), guanosine triphosphate (GTP, 10 mM; NEB, M2080S), and a custom biotinylated template-switching oligonucleotide (5′ Biotin-ATGGCUCGGAGAUGUGUAUAAGAGACAGUCUrGrG+G; Integrated DNA Technologies). Reagents were mixed immediately prior loading and kept on ice until droplet generation. Cells or nuclei were loaded into the 10x Genomics Chromium controller following the manufacturer’s protocol for 3′ v3.1 chemistry.

### cDNA amplification and ribosomal RNA depletion

Following reverse transcription, emulsion droplets were broken according to 10x recovery steps. cDNA was then treated with uracil-DNA glycosylase (UDG; NEB, M0280S) to remove dUTSO fragments at 37 °C for 50-60 min. This was followed by 5 cycles of amplification using 10x proprietary primers and a custom spike in primer (according to TotalX protocol, **Supplementary Note**). pre-amped cDNA (for 5 cycles) was cleaned up using 1.8X SPRI Select (Beckman Coulter, B23317) and cDNA corresponding to ribosomal RNA was depleted by Cas9-mediated cleavage using guide RNAs targeting mitochondrial and cytoplasmic ribosomal sequences (DASH). CRISPR ribonucleoprotein complexes were assembled from Alt-R S.p. Cas9 Nuclease V3 (NEB, 1081059), tracrRNA (IDT, 1072532), and a custom pool of 57 crRNAs (IDT) designed based on Smart-seq-total DASH oligos (**Supplementary Table 1**). The reaction was incubated at 37 °C for 60 min, followed by treatment with proteinase K (Thermo Fisher Scientific, EO0491) for 10 min at 56 °C. cDNA was then cleaned up using SPRI Select and amplified for additional 7-10 cycles using 10x proprietary primers and a custom spike in primer (see **Supplementary Note** for the detailed TotalX protocol).

### Library construction, size selection and sequencing

Amplified cDNA was separated into long- and short-fragment pools using SPRIselect magnetic beads (Beckman Coulter, Cat# B23318). To obtain the long-fragment pool, SPRIselect beads were added to amplified cDNA at a 0.6X bead-to-sample ratio, and bound fragments were retained according to the manufacturer’s instructions. Unbound fragments in the supernatant were further purified at a 1.2X bead ratio (1.8X final) to isolate the short-fragment pool. The long-fragment pool was processed according to the Chromium Single Cell 3′ v3.1 protocol (10x Genomics), using the Single Index Kit T Set A (PN-1000213/PN-2000240). Short fragments were indexed using custom primers through PCR amplification containing 10x Genomics 3′ v3.1 library amplification mix, SI primer and custom index primers (see **Supplementary Table 1**) For each 40 μl reaction, the following components were combined: 20 μl Amp Mix, 4 μl SI primer, 1 μl custom index primer (10 μM), 1 μl amplified cDNA (1 ng/μl), and 14 μl nuclease-free water. PCR conditions were as follows: 98 °C for 3 min; 12 cycles of 98 °C for 15 sec, 62 °C for 20 sec, and 72 °C for 2 min; and a final extension at 72 °C for 1 min. PCR products were purified with a 1.2X SPRIselect bead clean-up and eluted in 30 μl nuclease-free water. The resulting libraries were either split for further purification to remove very short fragments (below 200-250bp) and obtain the small RNA fraction, or used for optional miRNA enrichment. The miRNA(+) fragments were further size-selected from the short-fragment pool by enrichment of fragments corresponding to small RNAs (18–30 bp) using the Pippin Prep gel system (Sage Science, 3% agarose), following the manufacturer’s protocol. Short and long pools were recombined in desired ratios prior to final library quality control and sequencing. Resulting libraries were sequenced on a NovaSeq 6000 instrument (Illumina) with the following configuration: Read 1, 28 bp; Index 1, 8 bp; Index 2, 0 bp; Read 2, 91 bp.

### Data processing and alignment

Raw sequencing data were demultiplexed using Cell Ranger mkfastq (v8.0.1) and Read2 was trimmed using Cutadapt^52^ with the following parameters: -u 6 -a “AAAAAAAAAA;min_overlap=10” -m 18. Trimmed reads were then processed using cellranger count with a dual-pass alignment strategy to enable quantification of both long and short RNA species. Two custom reference transcriptomes were constructed from the GRCh38 primary assembly (Ensembl release 109; GRCh38.p13). Gene annotations were sourced from GENCODE v44 ^53^ and manually curated to include an expanded set of biotypes.

The primary (long RNA) reference included the following biotypes: protein_coding, lncRNA, miRNA, snRNA, snoRNA, scaRNA, tRNA, Mt_tRNA, vault_RNA, misc_RNA, antisense, scRNA, and immunoglobulin and T-cell receptor genes (e.g., IG_*, TR_*), along with their pseudogenes. tRNA annotations were appended using curated entries from GtRNAdb^54^. For viral infection studies, the dengue virus serotype 2 (DENV2) genome (GenBank NC_001474; GCF_000871845.1)^55^ and its corresponding annotation (modified_dengue.gtf) were appended.

The secondary (small RNA) reference was restricted to short RNA biotypes (miRNA, snoRNA, scaRNA) to enhance mapping specificity. tRNAs and viral transcripts were excluded from this reference.

Alignment, barcode correction, UMI collapsing, and gene quantification were performed using Cell Ranger’s internal rules, with modified alignment parameters specified in the cellranger-8.0.1/lib/bin/parameters.toml file to improve capture of short RNA species (≤18 bp). Key modified parameters included:

star_parameters = “--outFilterMismatchNoverLmax=0.05 --outFilterMatchNmin=18-- outFilterScoreMinOverLread=0 --outFilterMatchNminOverLread=0”

Many small non-coding transcripts—including miRNAs and other short RNAs—were found to be located within introns, exons, or overlapping lncRNAs (**Extended Fig. 1e**). In the standard Cell Ranger pipeline, reads overlapping multiple features are not counted, leading to underrepresentation of these transcripts. To address this, a two-step mapping and quantification strategy was employed.

Initial barcode calling and quality filtering were performed using the long RNA reference. Barcodes passing these filters were then used to extract matching data from the short RNA alignment. In accordance with the cellranger logic only reads containing the tag `xf:Z:25` in the BAM file were used for UMI counting and generation of the final gene expression matrix. The final gene expression matrix was constructed by substituting miRNA and other short RNA feature counts from the short-reference run into the long-reference matrix based on barcode concordance. Only barcodes confidently detected in the long RNA run were retained for downstream analysis.

Custom Python scripts were developed to merge the count matrices, resolve feature overlaps, ensure UMI count accuracy, and annotate transcript biotypes (see GitHub repository).

### Comparison to other methods

Publicly available HEK293T single-cell RNA-sequencing (scRNA-seq) datasets generated using the VASA-seq droplet-based protocol were downloaded from the NCBI Gene Expression Omnibus (GSE176588). For the 10x Genomics Chromium v3.1 dataset, data files were obtained directly from the 10x Genomics dataset page. To enable direct comparison, all datasets were first harmonized by mapping genes to Ensembl gene IDs, and only genes present in all datasets were retained for downstream analyses. Gene symbols were standardized by removing version suffixes (e.g., “.1”, “.2”), and gene detection rates were estimated based on unique, non-ambiguous gene symbols within each dataset.

For the TotalX dataset, to minimize the impact of putative doublets, barcodes corresponding to the top 8% of total UMI counts (reflecting the predicted doublet rate) were excluded. For datasets containing more than 1,000 cells, only cells with at least 5,000 detected UMIs were retained; smaller datasets were included in full. To compare gene detection rates and account for differences in sequencing depth between protocols, UMI counts for each cell were randomly downsampled to a fixed target of 20,000 UMIs per cell using a custom script. Cells with fewer than the target UMI count were excluded, except in datasets with fewer than 1,000 cells, where all cells were kept. Gene detection was assessed as the number of genes with at least one UMI detected per cell following downsampling. For comparisons of mean expression values between protocols, only genes shared across datasets were included.

### Doublet removal, cell-type annotation and downstream analysis

Doublets were removed using Scrublet^56^ with thresholds selected to match the expected doublet rate. Integration across brain samples was performed using scVI^57^ using protein-coding highly variable genes. Dimensionality reduction and clustering were conducted in Scanpy^58^ using the scVI latent space, followed by principal component analysis and UMAP^59^ for visualization. Differential gene expression analysis was performed using the Wilcoxon rank-sum test with Bonferroni correction. Non-coding RNA expression was quantified using log-normalized counts. Gene co-expression modules were identified using the Python implementation of weighted gene co-expression network analysis WGCNA^60^, with soft-thresholding and module detection as previously described.

For pseudotime analyses, developmental trajectories were inferred using PAGA and diffusion pseudotime in Scanpy. For further details refer to the GitHub page.

### Quantification of tRNA availability and demand

To estimate cell type–specific amino acid demand, we first obtained protein-coding sequences from GENCODE v44. For each protein-coding gene, the corresponding protein sequence was extracted, and the frequency of each standard amino acid was tabulated. Within each cell type, gene-level UMI counts were summed across all cells, yielding total expression for each gene per cell type. For genes with a mapped protein sequence, we calculated the total number of each amino acid required by multiplying the gene’s summed expression by the corresponding amino acid counts from the protein sequence. The aggregate for each amino acid per cell type thus reflects the overall translational demand for each amino acid. Amino acid demand matrices were stored as tables with amino acids as rows and cell types as columns. To quantify tRNA availability, we identified all tRNA genes present in the reference transcriptome (filtered for names containing the “tRNA” string). For each tRNA gene, the associated amino acid was inferred from the gene name, using regular expression parsing (e.g., extracting the amino acid label preceding “_tRNA”). We then summed UMI counts for all tRNA genes by cell type. For each amino acid, total tRNA expression was computed as the sum of all tRNA gene UMIs corresponding to that amino acid within each cell type. This yielded a matrix of tRNA “supply” per amino acid per cell type.

### Quantification of DENV2

DENV2 viral RNA abundance was quantified at the single-cell level using two complementary approaches. First, for each cell, the total viral RNA load was computed as the sum of normalized expression values for five DENV2-derived genes: *sfRNA1*, *sfRNA2*, *sfRNA3*, *sfRNA4*, and *POLY*. The resulting value (`viral_load`) was further normalized to the total cellular transcript count to calculate the percentage of viral transcripts per cell (`viral_pct`).

In parallel, a composite viral load score was generated using the `score_genes` function in Scanpy (v1.11.0), which computes the average expression of the specified viral genes after subtracting the average expression of a control gene set matched for overall expression distribution. This was performed for both the full viral gene set (`viral_load_score`, based on *sfRNA1–4* and *POLY*) and for the non-coding flaviviral RNAs alone (`sfrna_load`, based on *sfRNA1–4*).

These viral burden metrics—absolute, relative, and control-adjusted—were used to stratify cells for all downstream analyses, including response clustering and pathway enrichment. Unless otherwise specified, `viral_load` was used for most analyses.

Gene ontology and Reactome pathway enrichment analyses of differentially expressed genes were performed using the Python package GSEApy^61^ and g:Profiler^62^ respectively, applying a hypergeometric test with FDR correction to identify significantly enriched pathways among top-ranked genes.

### Correlation between miRNA and its targets

To assess the relationship between miRNA and target gene expression at single-cell resolution, only cells expressing the miRNA of interest were included in the analysis. For each cell, miRNA expression was quantified, and cells were partitioned into 30 quantile-based bins according to miRNA abundance using `pandas.qcut`. This approach enabled robust estimation of monotonic relationships between miRNA and target gene expression across the full dynamic range of miRNA levels observed.

Validated and predicted targets for each miRNA were compiled from TarBase^49^ and TargetScan 8.0^47^, respectively, and subsequently filtered to retain only those genes detected in the dataset. For each target gene, the mean expression within each miRNA expression bin was calculated. The Spearman rank correlation coefficient (ρ) between the binned mean miRNA expression and the corresponding target gene expression profile was then computed using the `scipy.stats.spearmanr` function. Only gene–miRNA pairs with sufficient coverage (at least five bins with non-missing values) were retained for further analysis. An analogous procedure was applied to all non-target genes to generate a background distribution of correlation coefficients, providing a reference for target specificity.

To elucidate the functional relevance of strongly anticorrelated miRNA targets, pathway enrichment analysis was performed. Validated target genes showing significant negative correlation (ρ < 0, adjusted P < 0.05) were subjected to pathway enrichment using the Reactome 2022 human gene set via the Enrichr interface (`gseapy` v0.10.8). Pathways with a false discovery rate (FDR) below 0.05 were considered significantly enriched. The top 20 enriched pathways were visualized as a scatterplot of –log (FDR) versus pathway name, with point size proportional to the odds ratio.

## Supporting information

Supplementary Note. TotalX Protocol

## Data availability

Processed data will be made available upon publication through [https://figshare.com/s/f6a2eea85980a6f7cd20]. Access to raw data may be requested and will be governed by a Data Use Agreement, which outlines the terms for academic use, citation, and redistribution.

## Code availability

Custom analysis scripts and instructions for Cell Ranger modifications are available via GitHub: https://github.com/alinaisakova/TotalX.

## Acknowledgments

This study was supported by the Chan Zuckerberg Biohub. A.I. was additionally supported by a Doc.Mobility Fellowship from the Swiss National Science Foundation. SE is a Chan Zuckerberg Biohub San Francisco Investigator who is also supported by National Institute of Allergy and Infectious Diseases grants RO1AI158569, Defense Threat Reduction Fundamental Research to Counter Weapons of Mass Destruction grant HDTRA11810039, and Investigator Initiated Awards W81XWH1910235 and W81XWH2210283 as well as Expansion Award W81XWH2110456 from the Department of Defense office of the Congressionally Directed Medical Research Programs/Peer Reviewed Medical Research Program. Finally, we are deeply grateful to the donors and their families for their invaluable contributions to this research. Human brain tissue used in this study was obtained through the coordinated efforts of the brain research neurosurgery teams at Stanford University. We thank these extended teams for their expertise, dedication, and care in tissue acquisition.

## Author Contributions

A.I. and S.R.Q. conceptualized the study. A.I. designed the study. A.I. and I.C. performed HEK293T and PBMC TotalX experiments. S.S. performed Huh7 and Huh7-DENV2 cell culture. A.I., D.D.L., A.E.E. and R.S. performed brain tissue processing. A.I. performed TotalX on the brain samples. A.D and N.N. sequenced TotalX samples. A.I. designed the computational pipeline and performed data analysis. A.I. wrote the manuscript with direct input from D.D.L., S.S, S.E., and S.R.Q. All authors reviewed and approved the final manuscript.

## Supplementary Information

Supplementary Table 1. Custom oligonucleotides used in this study.

Supplementary Table 2. Differentially expressed genes (DEGs) identified in the PBMC dataset (adjusted P < 0.05).

Supplementary Table 3. Gene modules identified in the PBMC dataset, comprising both coding and non-coding genes.

Supplementary Table 4. Differentially expressed genes in Huh7-DENV2: Quiescent_highVL vs. Quiescent_lowVL.

Supplementary Note. TotalX protocol.

## EXTENDED FIGURES

**Extended Figure 1.**
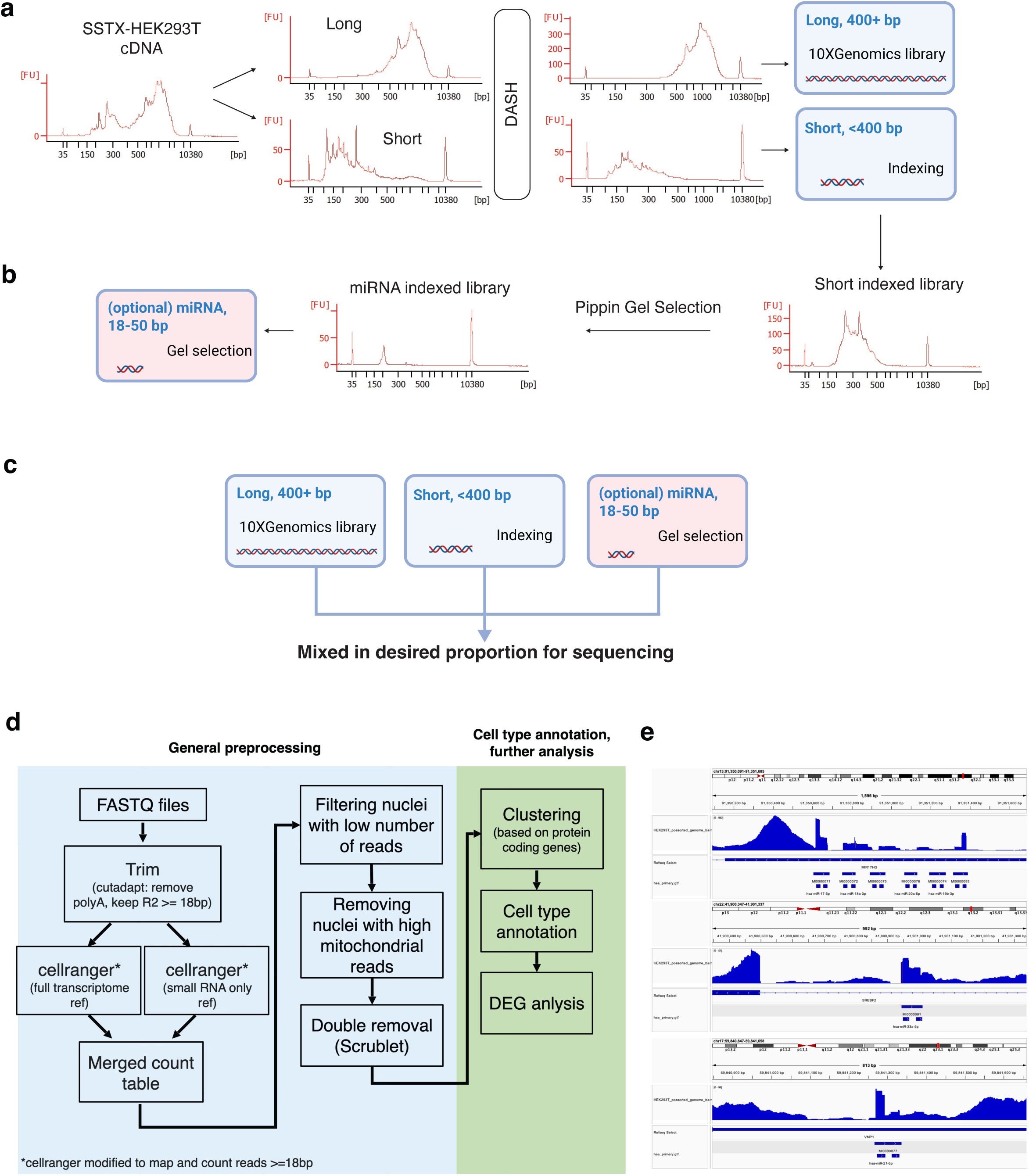
TotalX experimental workflow and analysis pipeline. **a.** Schematic overview of the TotalX workflow. Total RNA is enzymatically polyadenylated and reverse transcribed using a custom template-switching oligo (TSO). The TSO is then selectively removed, followed by Cas9-based rRNA depletion (DASH) at the cDNA level. Resulting cDNA fragments are size-separated, with short (<400 bp) and long (>400 bp) fractions indexed independently and pooled for sequencing. An optional gel-purified miRNA fraction (~18–50 bp) enables targeted enrichment of mature small RNAs. **b.** Structure of the short-fragment indexed library. The short-fragment indexed libraries can optionally undergo Pippin gel selection system to obtain miRNA-enriched libraries. **c.** Workflow for combining short RNA, miRNA-enriched and standard cDNA fractions. The resulting composite library captures a broad range of transcript sizes and biotypes. **d.** Cell Ranger-based data analysis pipeline of TotalX. **e.** Examples of gene coverage profiles for representative miRNAs located within the lncRNA locus (top), intron of a protein coding gene (middle) and exon of a protein coding gene (bottom). Bam file used for the coverage plots was generated with the modified Cell Ranger pipeline.

**Extended Figure 2.**
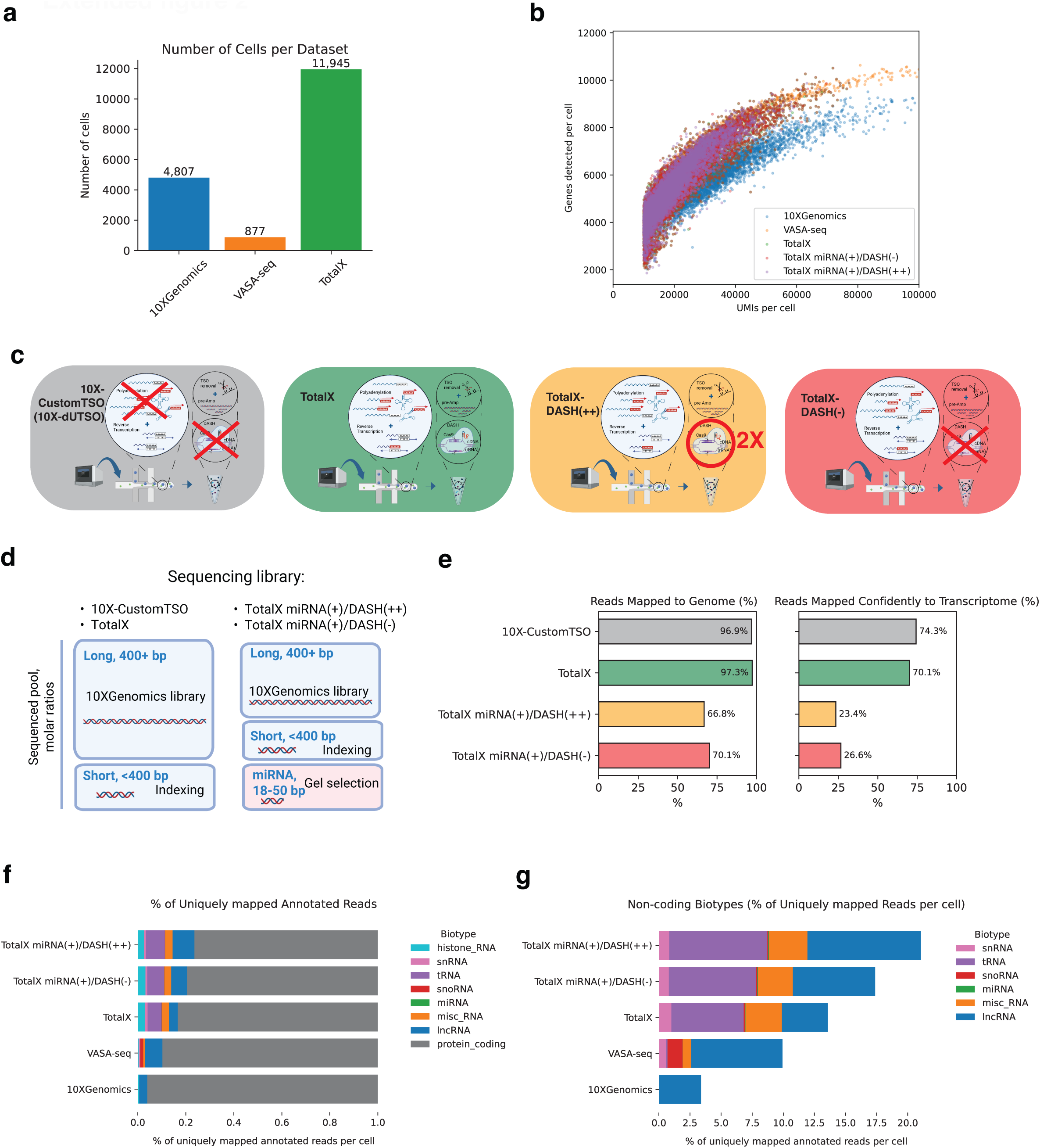
Performance benchmarking of TotalX libraries by gene complexity, biotype diversity, and throughput. **a.** Number of cells per dataset for 10x Genomics, VASA-seq, and TotalX libraries. **b.** Relationship between UMIs per cell and number of genes detected per cell across datasets and library configurations. **c.** Schematic overview of library preparation workflows, including 10x-CustomTSO (10x-dUTSO), TotalX, and TotalX with or without DASH treatment. Three representative configurations are illustrated: TotalX with miRNA enrichment and two rounds of DASH (miRNA(+)/DASH(++)), TotalX with miRNA enrichment but no DASH (miRNA(+)/DASH(–)), and standard TotalX without additional modifications. **d.** Summary of sequencing library construction and pooling strategies, for each tested library configuration. **e.** Percentage of reads mapped to the genome and confidently mapped to the transcriptome across different library types. **f.** Fraction of uniquely mapped annotated reads per cell, stratified by RNA biotype. **g.** Contribution of non-coding RNA biotypes (e.g., snRNA, tRNA, snoRNA, miRNA, miscRNA, lncRNA) to uniquely mapped reads per cell across methods.

**Extended Figure 3.**
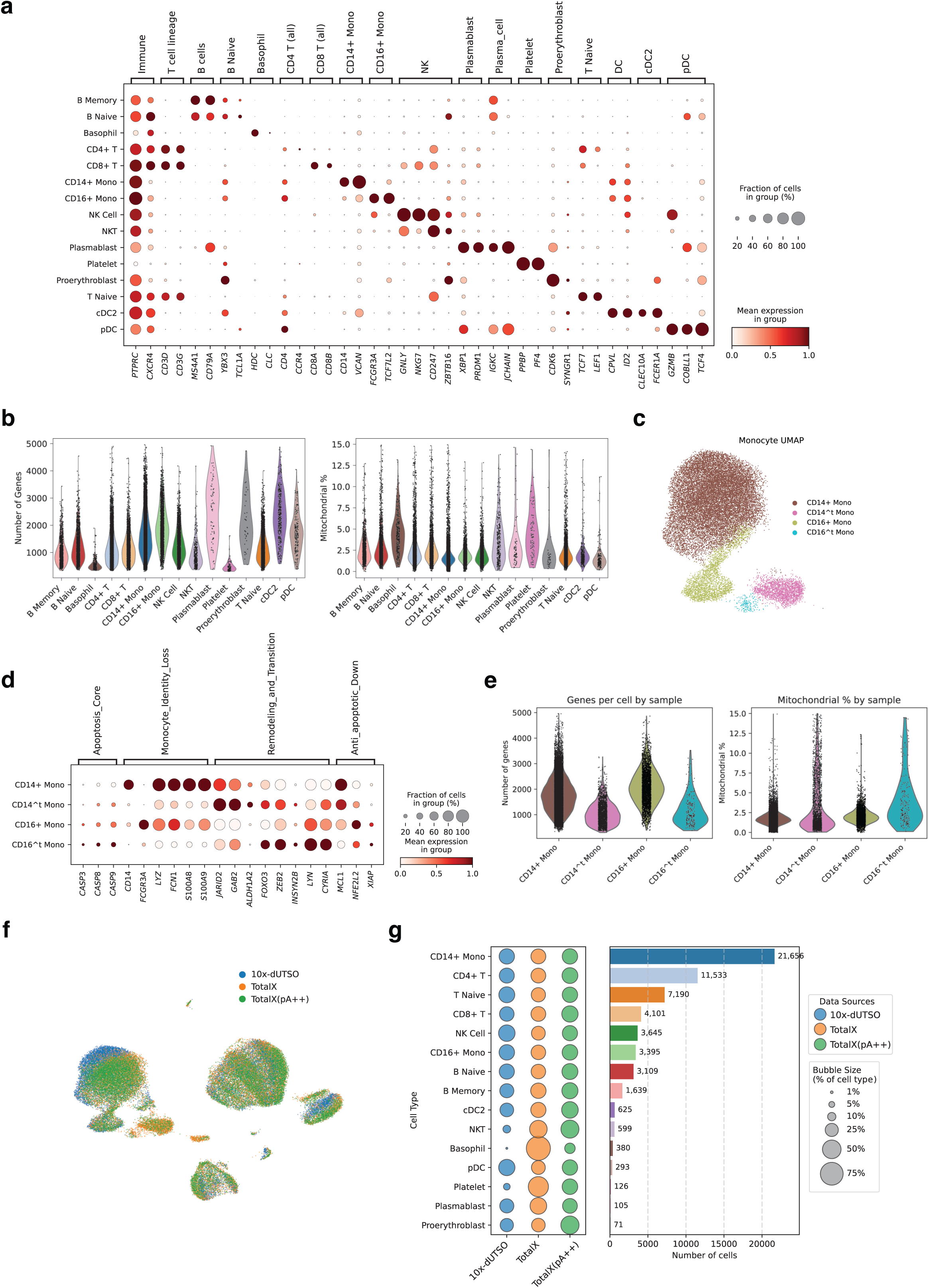
Comparative profiling of PBMC subtypes across protocols and sensitivity of monocytes to enzymatic polyadenylation. **a.** Dot plot showing expression of canonical marker genes across annotated immune cell types, including major T cell subsets, monocytes, NK cells, dendritic cells, basophils, plasmablasts, and platelets. Marker genes were used for manual annotation of cell clusters in TotalX, 10x-dUTSO, and TotalX+miRNA(+) protocols. **b.** UMAP of PBMCs colored by protocol, demonstrating that TotalX and TotalX + retain consistent clustering and recover all expected immune populations observed in the 10x-dUTSO control. **c.** Left: Bubble plot of relative frequencies of immune cell types across protocols. Right: Absolute numbers of cells recovered by TotalX, TotalX pA++ and 10x-dUTSO. TotalX recovers comparable or greater numbers of low-RNA-content populations (e.g., platelets, basophils, proerythroblasts). **d.** Violin plots showing the number of genes (left) and UMIs (right) detected per cell type. Low-content cell types such as platelets and basophils show improved gene detection under TotalX, while other major cell types are comparable across protocols. **e.** UMAP focused on monocyte populations reveals a distinct subcluster of CD14+ monocytes in TotalX + showing transcriptional divergence, indicative of sensitivity to protocol conditions. **f.** Dot plot showing selective upregulation of JARID2, GAB2, and APOBEC3A, along with downregulation of canonical monocyte markers (e.g., LYZ, CD14), in CD14+ monocytes exposed to increasing concentrations of ATP and polyA polymerase. These cells also show enrichment for apoptosis and chromatin remodeling genes. **g.** Violin plots showing global decrease in gene detection and increased mitochondrial RNA content in the sensitive monocyte subpopulation, consistent with stress or partial identity loss under certain enzymatic conditions.

**Extended Figure 4.**
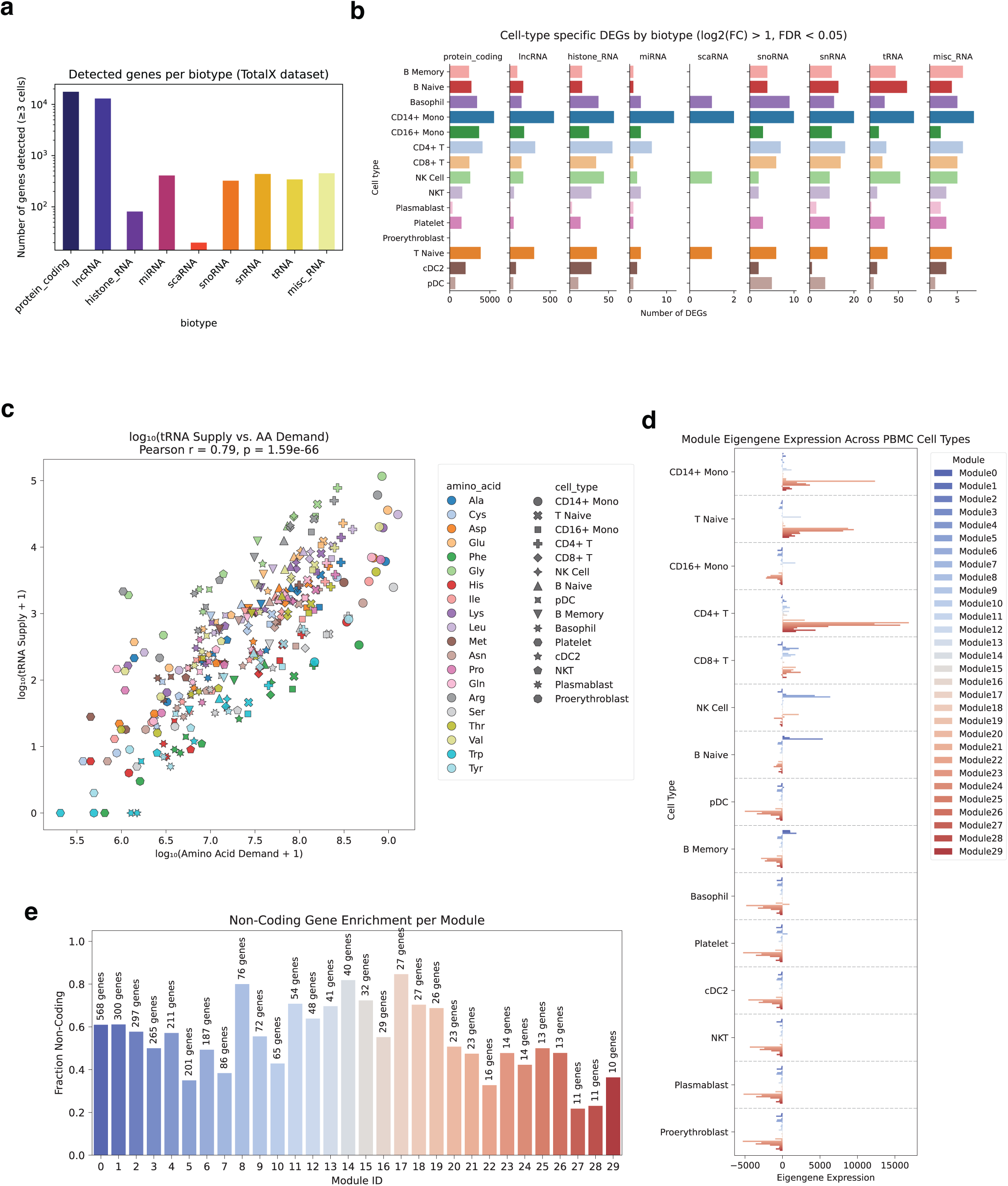
Cell-type specificity and co-expression patterns of non-coding RNAs in PBMCs. **a.** Number of genes detected per RNA biotype (minimum expression in ≥3 cells) in the TotalX PBMC dataset. Biotypes include protein-coding genes, lncRNAs, tRNAs, snoRNAs, snRNAs, histone RNAs, miRNAs, and miscellaneous RNAs, reflecting TotalX’s capacity to capture broad transcript classes. **b.** Cell-type specific differentially expressed genes (DEGs) stratified by RNA biotype. Bars indicate the number of DEGs (FDR < 0.01) for each cell type and RNA class. While protein-coding genes remain dominant, multiple non-coding biotypes exhibit strong cell-type–specific expression—e.g., lncRNAs in monocytes, miRNAs in lymphocytes, and snRNAs in plasmacytoid dendritic cells (pDCs). **c.** Relationship between amino acid demand (based on codon usage of expressed genes) and tRNA supply (measured by TotalX) across immune cell types. The strong correlation (Pearson r = 0.79, p = 1.59e–66) confirms internal consistency and biological relevance of captured tRNA profiles. Selected amino acids such as Arginine and Glycine show relative oversupply, whereas Tryptophan and Phenylalanine are undersupplied. **d.** Eigengene expression of co-expression modules across PBMC cell types, as identified by WGCNA. Blue modules are enriched in protein-coding content, whereas red modules include higher fractions of non-coding RNAs. Module 24 is enriched in CD14+ monocytes and T cells, while Module 7 is platelet-specific. **e.** Fraction of non-coding genes in each WGCNA module. Several modules (e.g., 1, 2, 6, 7) include substantial non-coding content, suggesting that cell identity and function are encoded in mixed coding/non-coding programs. Total gene counts per module are indicated above each bar.

**Extended Figure 5.**
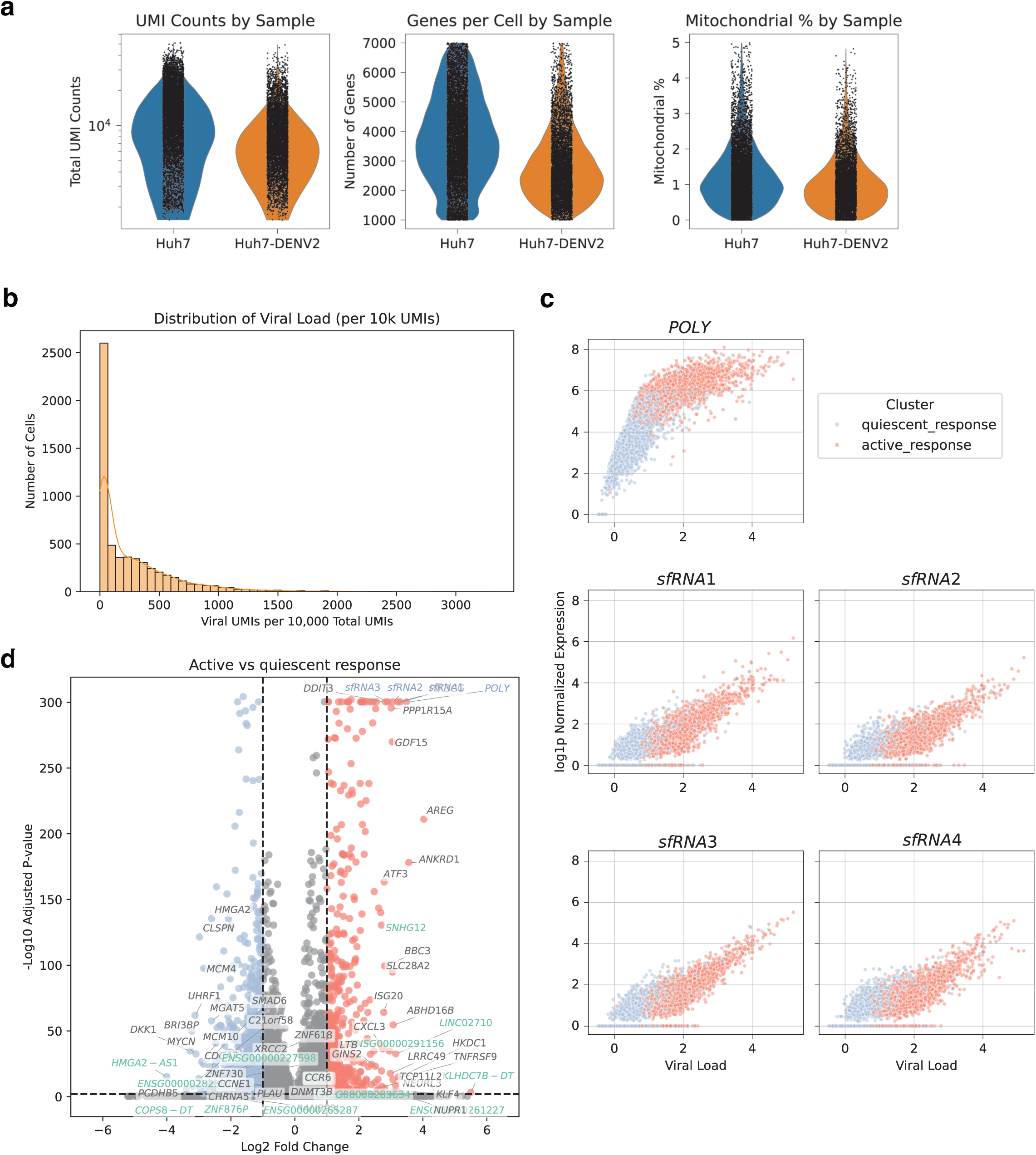
Sensitivity of TotalX for viral transcript detection and host response stratification in DENV2-infected cells. **a.** Quality control metrics comparing uninfected (Huh7) and DENV2-infected (Huh7-DENV2) samples. Violin plots show total UMI counts, number of detected genes, and mitochondrial transcript percentage per cell. **b.** Histogram showing the distribution of viral load per cell, defined as dengue virus–derived UMIs per 10,000 total UMIs. **c.** Scatter plots showing normalized expression of individual viral transcripts (*POLY, sfRNA1–4*) versus viral load per cell. Expression of both structural (*POLY*) and non-coding (*sfRNAs*) viral RNAs scales with total viral burden. Cells are colored by transcriptional state: quiescent response (blue) and active response (red). **d.** Volcano plot comparing gene expression between active and silent infection states. Several transcription factors (e.g., *ATF3*, *DDIT3*) and immune response genes are upregulated in active response, while chromatin remodeling factors and proliferation-associated genes are enriched in the silent state. Gene names are colored as follows: coding in gray, non-coding in green and viral in blue.

**Extended Figure 6.**
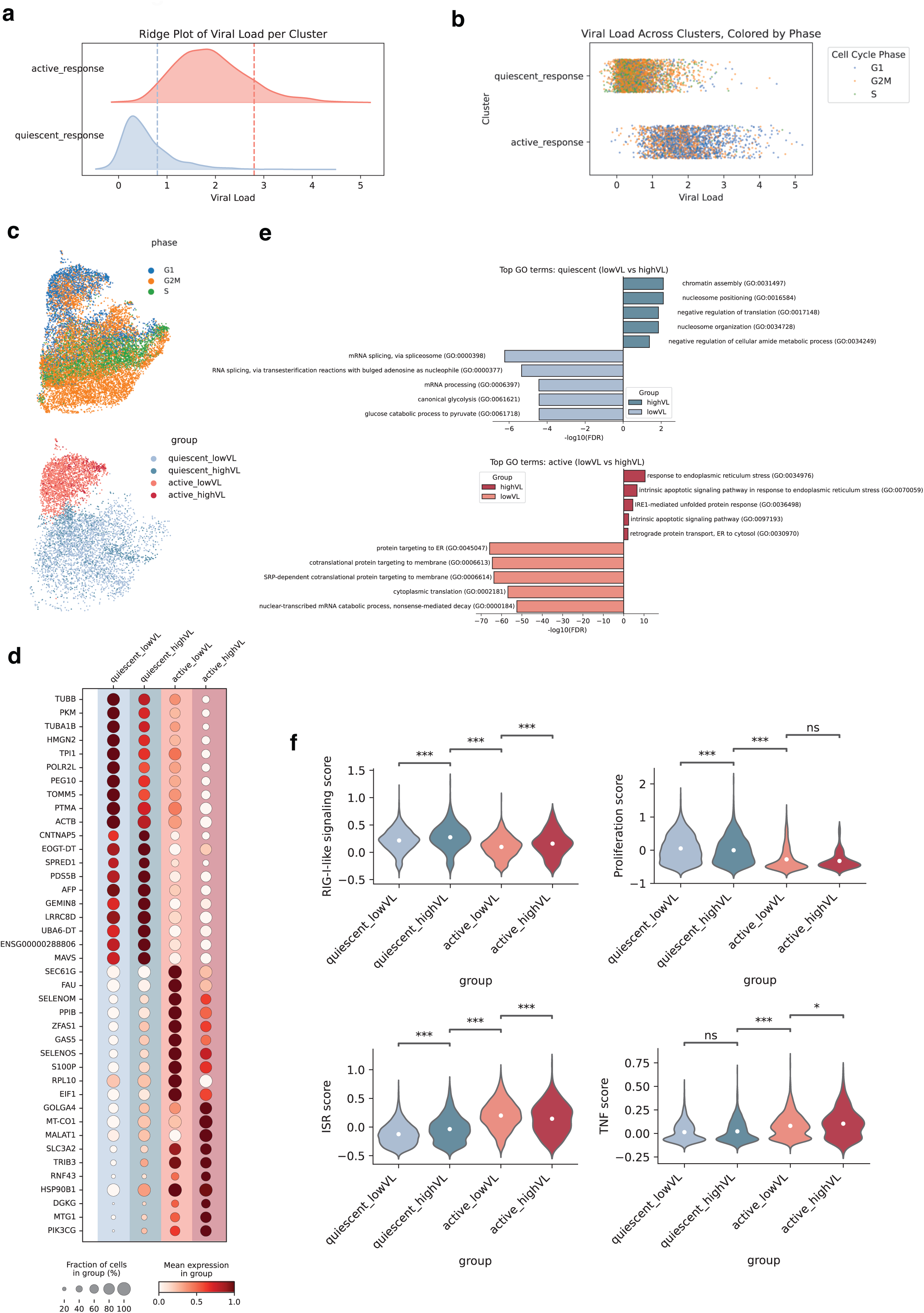
Viral load–based stratification of host response in DENV2-infected cells. **a.** Ridge plot showing the distribution of viral RNA levels across transcriptional clusters. Cells are grouped into two major infection states—quiescent infection (blue) and active infection (red)—based on host transcriptomic profiles. Although overall viral load can be similar between states, active cells display a robust host response, while quiescent cells remain largely unresponsive. **b.** Viral load across infection groups, colored by cell cycle phase (G1, S, G2/M). **c.** UMAP embeddings of infected cells colored by cell cycle phase (top) and infection group (bottom): quiescent_lowVL, quiescent_highVL, active_lowVL, active_highVL. **d.** Gene expression across four states. Cells were stratified into quiescent/active × low/high viral load groups. Dot plot shows expression patterns of differentially expressed genes. **e.** GO enrichment analysis for genes differentially expressed between high and low viral load cells within each infection state. Top: In quiescent response, high-load cells are enriched for chromatin remodeling, RNA splicing, and nucleosome-related processes. Bottom: In active response, high-load cells show enrichment for unfolded protein response (UPR), ER stress, and protein targeting to membranes. **f.** Violin plots showing module scores for key biological processes across the four infection groups.

**Extended Figure 7.**
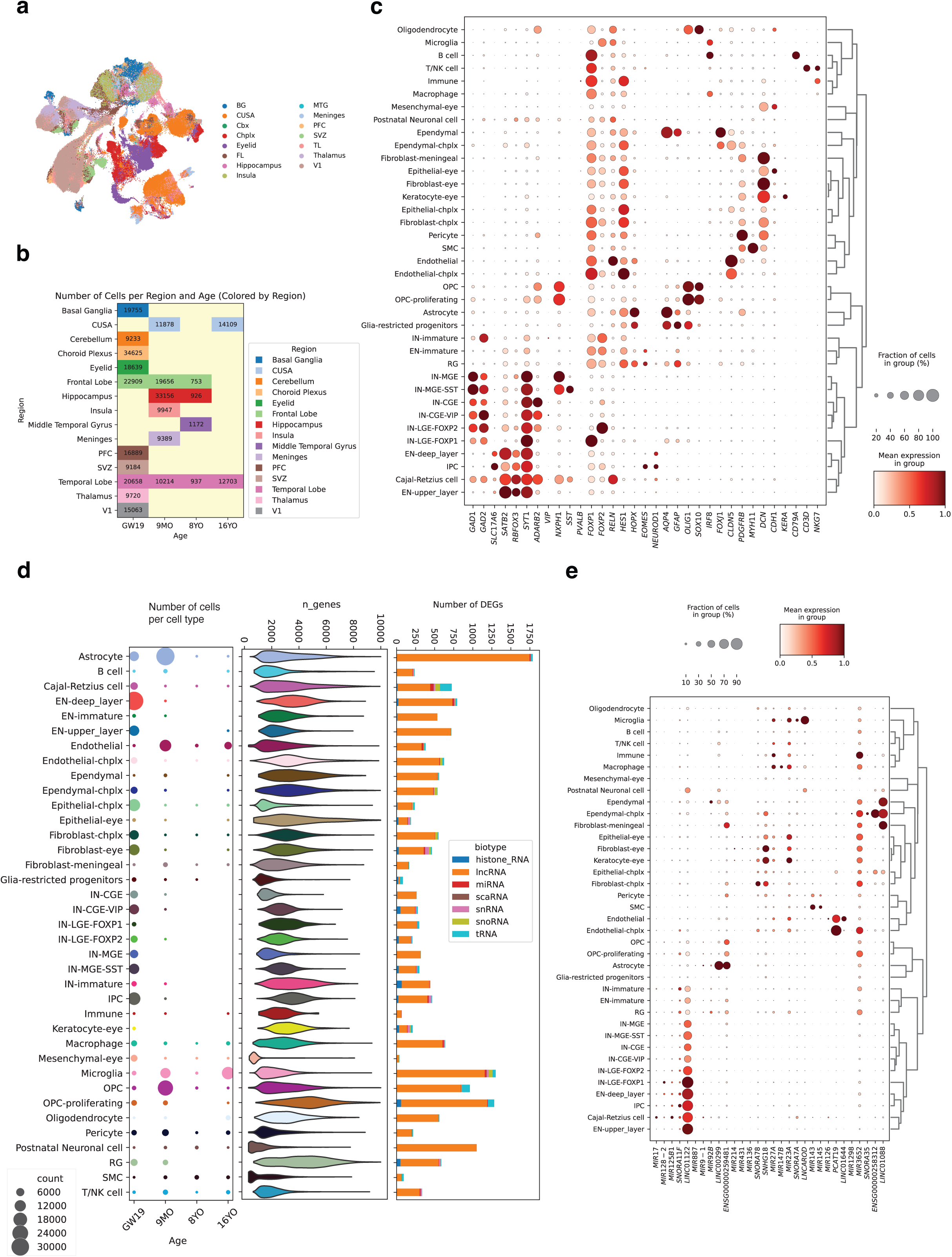
Regional coverage, cell-type diversity, and age-dependent non-coding RNA expression in the developing human brain. **a.** UMAP embedding of 301,515 single cells profiled using TotalX, colored by brain region of origin. The dataset includes cortical and subcortical structures, non-parenchymal compartments (e.g., meninges, choroid plexus), and ocular-associated tissues (e.g., optic region, eyelid), reflecting broad anatomical sampling across developmental stages. **b.** Number of cells recovered from each brain region across four developmental timepoints: GW19, 9MO, 8YO, and 16YO. Bars are color-coded by region and capture both prenatal and postnatal contributions to the dataset. **c.** Dot plot showing marker gene expression across annotated cell types. Cell types span neuronal, glial, immune, and non-neural lineages and are hierarchically clustered by expression patterns. **d.** Cell-type–resolved gene detection across development. **Left:** Dot plot showing the number of cells per annotated cell type at each age. **Center:** Violin plots depicting the number of detected genes per cell. **Right:** Bar plots showing the number of differentially expressed non-coding RNAs per cell type and RNA biotype (Wilcoxon test, log fold change > 1, FDR < 0.01). Histone RNAs are enriched in proliferating progenitors, while other biotypes vary by lineage and developmental stage. **e. Selected cell-type–specific non-coding RNA markers.** Dot plots display expression of non-coding RNAs across brain cell types, grouped by broad cell classes. Same as in figure 4d.

**Extended Figure 8.**
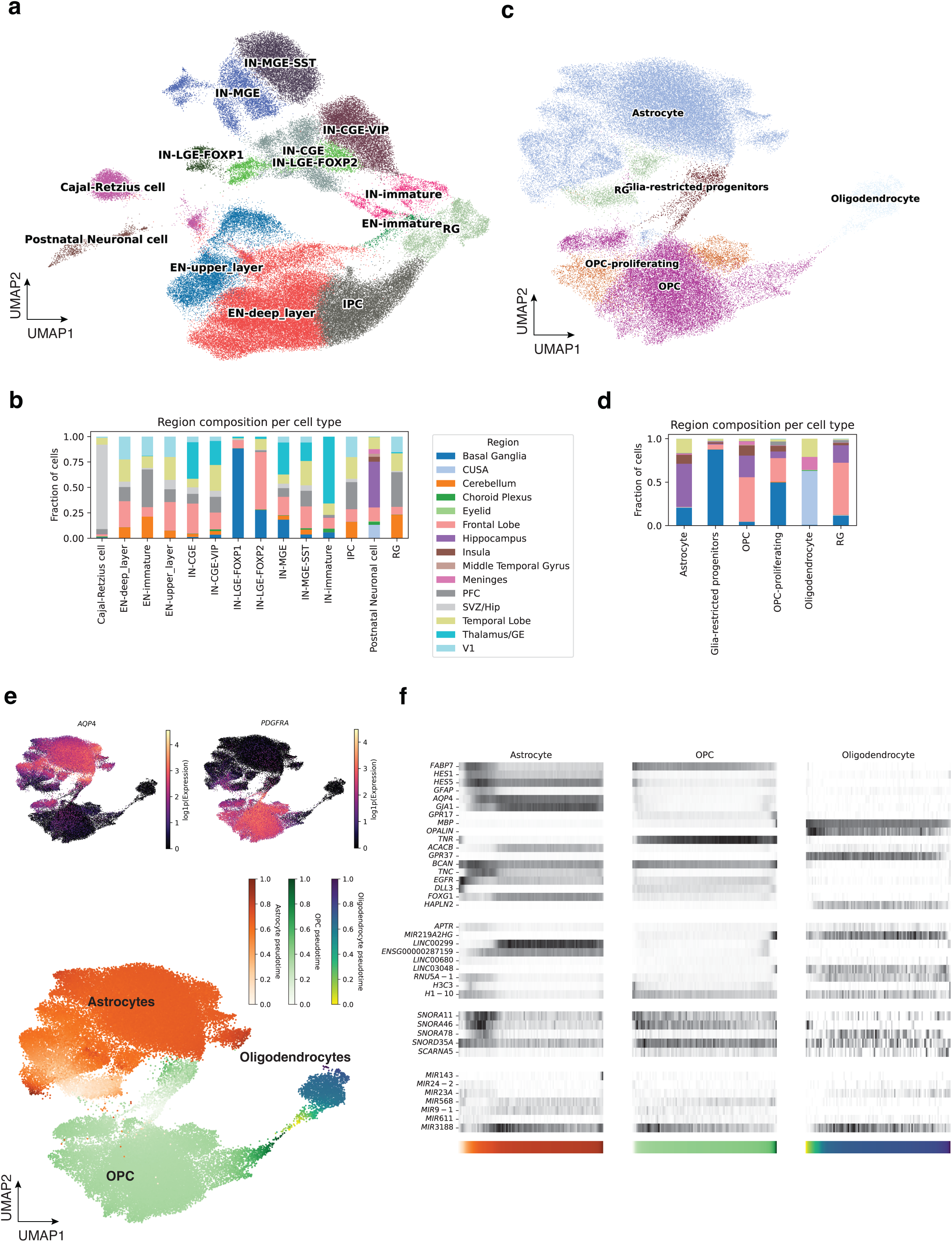
Regional and molecular diversity of glial and neuronal subtypes in the developing human brain. **a.** UMAP embedding of excitatory and inhibitory neuron subtypes, colored by cell type. Major cell types include Cajal-Retzius cells, postnatal neurons, deep and upper-layer excitatory neurons (EN), interneurons from MGE, CGE, and LGE origins (e.g., IN-MGE-SST, IN-CGE-VIP, IN-LGE-FOXP1), and radial glia (RG), reflecting broad neuronal lineage heterogeneity. **b.** Bar plot showing regional contributions to each neuronal subtype. Bars represent the fraction of cells per region for each annotated neuronal cell type. **c.** UMAP embedding of glial subtypes, including astrocytes, oligodendrocytes, OPCs, OPC-proliferating cells, and RG-like progenitors. **d.** Bar plot showing regional contributions to each glial cell type. As in (b), bars indicate the fraction of cells derived from each brain region. **e.** Diffusion map of the glial lineage, showing trajectories from RG to mature OPCs, astrocytes, and oligodendrocytes. **Top:** Expression gradients of canonical markers (e.g., *AQP4*, *PDGFRA*) across the lineage. **Bottom:** Pseudotime-colored embedding with annotations of cell type transitions. **f.** Heatmaps showing expression of selected coding and non-coding RNA markers across astrocytes, OPCs, and oligodendrocytes that change along the pseudotime trajectory. Rows represent genes; columns represent cells ordered by pseudotime (100 bins). Color bars denote pseudotime, consistent with panel (e).

**Extended Figure 9.**
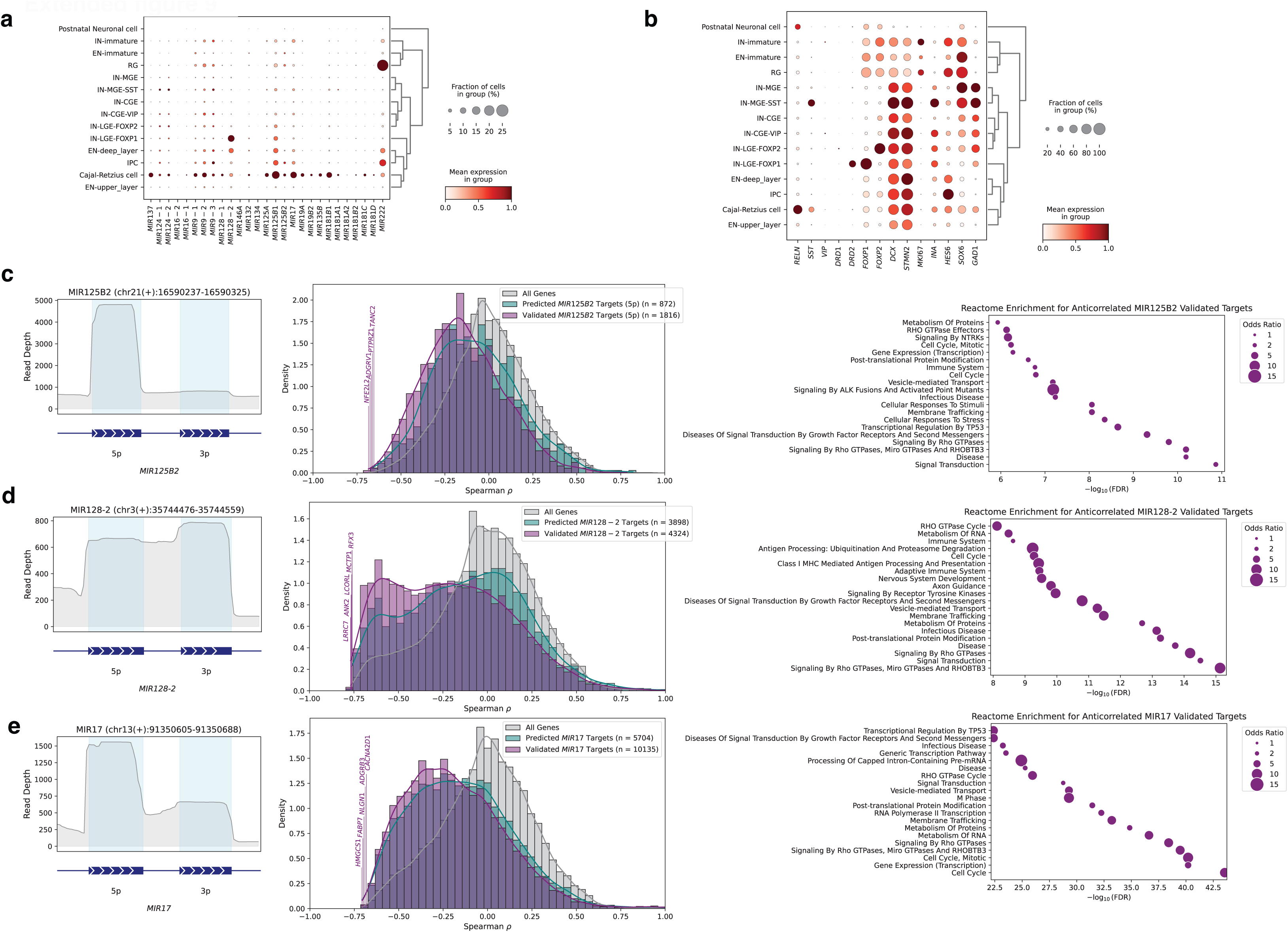
Cell-type–specific expression and predicted target relationships for non-coding RNAs in developing neurons. **a.** Dot plot showing the cell-type–specific expression of selected miRNAs across neuronal cell types. **b.** Dot plot showing the expression of selected mRNAs marker genes across neuronal subtypes, using the same format as in (a). **c–e.** Genomic loci, correlation distributions, and pathway enrichments for representative miRNAs with strong cell-type–specific expression and anticorrelated targets. Same as in Figure 5. **c.** *MIR125B2*. **Left:** Genomic coordinates and coverage profile. **Center:** Distribution of Spearman correlation coefficients between *MIR125B2* expression and predicted or validated target gene expression across cells. **Right:** Reactome pathway enrichment analysis of anticorrelated validated targets (odds ratio is represented by dot size). **d.** *MIR1262*; panels follow the same format as in c. **e.** *MIR17*; panels follow the same format as in c.

