## Supplementary Note. TotalX Protocol for "Beyond polyA: scalable single-cell total RNA-seq unifies coding and non-coding transcriptomics"

### TotalX on 10x Genomics Single Cell 3’ v3.1 with Custom TSO and RT Amplification Oligos

#### Step 1.1: Incorporation of Custom Oligonucleotides

**Custom Oligos:** - **Template Switch Oligo (TSO)** (from IDT, PAGE purified):
Biotin-ATGGCUCGGAGAUGUGUAUAAGAGACAGUCUrGrG+G

**RT Amplification Oligo (D01-RT-AMP-short):**
GCTCGGAGATGTGTATAAGAGACAG

**Instructions:**
- Replace the standard 10x Genomics TSO (PN-3000228) with the custom TSO above in the RT Master Mix.
- Add D01-RT-AMP-short oligo during cDNA amplification steps as detailed below.

#### Step 1.2: GEM Generation & Barcoding

- Follow the standard 10x Genomics protocol for chip loading, GEM generation, and GEM-RT incubation, using the modified RT master mix.

**Table 1. Custom Reverse Transcription (RT) Reaction Mix**

| Component | Volume (μl) | Notes/Source |
| --- | --- | --- |
| RT reagent B | 18.8 | 10x Genomics |
| Reducing agent B | 2.0 | 10x Genomics |
| RT enzyme C | 8.7 | 10x Genomics |
| dUTSO (custom template switch oligo) | 2.4 | Custom, replaces standard TSO |
| PolyA polymerase (5 U/μl) | 2.0 | NEB, #M0276S |
| ATP (10 mM) | 1.0 | NEB, #M0276S |
| Vaccinia capping enzyme (10 U/ul) | 1.0 | NEB, #M2080S |
| SAM (2 mM) | 0.5 | NEB, #M2080S |
| GTP (10 mM) | 0.5 | NEB, #M2080S |
| dH₂O | 0.1 |  |
| **Total RT Master Mix** | **37.0** |  |

- Mix 37 μl RT master mix with 38 μl cell suspension for 75 μl total volume.

#### Step 2: Post GEM-RT Cleanup & cDNA Amplification

##### 2.1: Post GEM-RT Cleanup (Dynabeads)

- Follow standard protocol for Recovery Agent addition, Dynabeads cleanup, and elution solution preparation.
- **Elute with 20.5 μl Elution Solution I** (modified volume), transfer 20 μl to a new tube strip.

##### 2.2: UDG Digestion

- Add **5 μl UDG** and **2.8 μl UDG buffer (10X)** (NEB, # M0280S) to the eluted 20 μl of cDNA.
- Incubate at 37°C for 50–60 minutes.

##### 2.3: First cDNA Amplification with Custom Oligos (“1st pre-AMP”)

- On ice, prepare the following for each sample:

| Component | Volume (μl) |
| --- | --- |
| UDG-digested cDNA | 28 |
| dH₂O | 7 |
| Amp Mix (10x Genomics) | 50 |
| cDNA primer (10x Genomics) | 15 |
| **D01-RT-AMP-short** (100 μM) | 0.5 |
| **Total Volume** | **100** |

- **Thermal cycling:**
  - 98°C for 3:00 min
  - 5 cycles:
    - 98°C for 15 sec
    - 62°C for 20 sec
    - 72°C for 2:00 min
  - Final extension: 72°C for 1:00 min
  - Hold at 4°C
  - Use SPRI Select to clean up the cDNA with 1.8X bead to sample ratio.

##### 2.4: DASH (Depletion of Abundant Sequences by Hybridization)

- **Guide RNA Preparation:**
  - Resuspend each crRNA oligo (IDT) to 100 μM in IDTE buffer.
  - Pool crRNAs (2 μl from guides 1–31, 1 μl from 32–57) for 88 μl total.
  - Resuspend each tracrRNA to 100 μM.
  - Mix pooled crRNA and tracrRNA equimolar to make 10 μM duplex.
  - Heat duplex at 95°C for 5 min, cool to room temp, store aliquots at –80°C.
- **RNP Complex Assembly:**
  - 10 μl 10 μM crRNA:tracrRNA duplex
  - 1 μl S.p. Cas9 (20 μM, NEB, #M0386S)
  - 5 μl 3.1 NEB buffer (10X)
  - 29 μl dH2O
  - Incubate at room temp for 5–10 min.
- **In vitro Digestion:**
  - 10 μl 3.1 NEB buffer (10X)
  - 21 μl 1 μM Cas9 RNP
  - 20 μl pre-amped cleaned-up cDNA
  - 49 μl dH20
  - 100 μl total, incubate at 37°C for 60 min.
  - Add 1 μl Proteinase K (20 mg/mL), incubate 56°C for 10 min.
  - Clean-up using 1.8X SPRI Select; elute in 28 μl.

##### 2.5: Second cDNA Amplification with Custom Oligos (“2nd pre-AMP”)

- On ice, for each sample:

| Component | Volume (μl) |
| --- | --- |
| cDNA (from above) | 28 |
| dH₂O | 7 |
| Amp Mix (10x Genomics) | 50 |
| cDNA primer (10x Genomics) | 15 |
| **D01-RT-AMP-short** (100 μM) | 0.25 |
| **Total Volume** | **100** |

- **Thermal cycling:**
  - 98°C for 3:00 min
  - 7 cycles:
    - 98°C for 15 sec
    - 62°C for 20 sec
    - 72°C for 2:00 min
  - Final extension: 72°C for 1:00 min
  - Hold at 4°C
- **SPRIselect Clean-up:**
  - **1st Bead Selection to purify Large Fragments (500-2000+bp):**
    Add 60 µl (0.6X) resuspended SPRI Select beads to 100 µl amplified DNA solution (this will keep fragments larger than 500 bp on the beads). Mix well by pipetting up and down at least 10 times. Incubate for 5 minutes at room temperature. Place the tube on a magnetic rack to separate the beads from the supernatant. After the solution is clear (about 5 minutes), carefully transfer the supernatant to a new tube **(Caution: do not discard the supernatant).** Keep beads, that contain the large fragments, on the magnet. Wash two times with 80% EtOH. Elute in EB.
  - **2nd Bead Selection to purify small Fragments (200- 500bp):**
    This step will bind the desired fragment sizes (contained in the **saved supernatant** from Step 1) to the beads. Add 140 µl (1.2X, will make 1.8X total) resuspended SPRI Select beads to the supernatant, mix well and incubate for 5 minutes at room temperature. Put the tube on a magnetic rack to separate beads from supernatant. After the solution is clear (approximately 3 minutes), carefully remove and discard the supernatant. Be careful not to disturb the beads that contain DNA targets **(Caution: do not discard beads)**. Wash two times with 80% freshly prepared ethanol. Elute in EB.

#### Step 3: Indexing Amplification of short fragments

| Component | Volume (μl) |
| --- | --- |
| Amp Mix | 20 |
| SI primer | 4 |
| Custom index primer (10 μM) | 1 |
| Amplified cDNA (1 ng/μl) | 1 |
| dH₂O | 14 |
| **Total** | **40** |

- PCR:
  - 98°C for 3 min
  - 98°C for 15 sec
  - 62°C for 20 sec
  - 72°C for 2 min
  - 72°C for 1 min
- SPRI Select clean-up: 1.2X, elute in 30 μl.
- Potentially split: 1) use for Pippin gel for miRNA (+) fraction and 2) perform additional clean-up to remove very short fragments and obtain small RNA fraction.

**All other steps** (library QC, quantification, sequencing) proceed per the standard 10x Genomics protocol unless otherwise specified.
